# An all-atom view into the disordered interaction interface of the TRIM5*α* PRYSPRY domain and the HIV capsid

**DOI:** 10.1101/2024.12.06.627233

**Authors:** Liam Haas-Neill, Deniz Meneksedag-Erol, Ibraheem Andeejani, Serena del Banco, Jeannette Tenthorey, Sarah Rauscher

## Abstract

Tripartite motif-containing protein 5, alpha isoform (TRIM5*α*) is an innate immune factor that provides rhesus macaques with immunity to HIV. Despite high sequence similarity to rhesus TRIM5*α*, human TRIM5*α* weakly restricts HIV without the introduction of mutations. The structural underpinnings of this functional difference are poorly understood because the interaction interface between TRIM5*α* and its target, the HIV capsid, involves intrinsically disordered regions. Here, we use all-atom molecular dynamics simulations to study several TRIM5*α* variants: rhesus TRIM5*α*, human TRIM5*α*, and human TRIM5*α* with an R332P mutation (a mutation known to enhance HIV restriction). Our data reveal differences in the conformational ensembles of wild-type and R332P human TRIM5*α*, including a significant increase in the formation of a turn that includes the mutation site residue. We also carried out simulations of rhesus TRIM5*α* in complex with the HIV capsid protein. Our results indicate that the variable loops of TRIM5*α* are highly flexible, both in solution and in the complex with the capsid, indicative of a fuzzy interaction interface. Simulations of the complex, as well as experimental infections, indicate the basis for weak HIV restriction by human TRIM5*α* that is improved by an R332P mutation: replacing the positively charged arginine with a neutral residue decreases electrostatic repulsion with residue R82 of the capsid protein, which can be phenocopied by neutralizing mutations at capsid residue R82. Overall, our simulations provide a first view of the atomistic details of the HIV capsid-TRIM5*α* binding interface and a molecular mechanism for the observed functional differences between variants of the TRIM5*α* protein.

**Significance Statement:** Humans and viruses are engaged in an evolutionary arms race, which involves evolutionary cycles in which viruses escape host defense and hosts evolve to chase after them. The protein TRIM5*α* is a crucial part of the immune system’s defense against viruses, and the region of TRIM5*α* that interacts with viral capsids is evolving rapidly. While a wealth of structural information is separately available for the HIV capsid protein and TRIM5*α*, the interaction interface between these two proteins remains poorly understood because it involves multiple disordered regions. We present a first atomistic view of the TRIM5*α*-capsid interaction interface using all-atom molecular dynamics simulations. We use our model of the interface and the probabilistic description of interactions to explain the improved HIV restriction of a mutation in the human TRIM5*α* protein, which has been proposed for human gene therapy, in terms of specific electrostatic interactions. Finally, we validate our findings with experimental infections. Our data provide detailed structural insight into how TRIM5 evolves improved restriction against its retroviral targets.

## Introduction

Viruses and their hosts are in an ongoing evolutionary battle in which both species constantly accumulate advantageous mutations to survive. These mutations accumulate at sites of direct interaction and presumably modulate changes in affinity between host and viral proteins, to the advantage of one side or the other.^1^ Host restriction factors are an important front-line defence mechanism against viruses in this evolutionary arms race.^2^ Restriction factors are host proteins that counteract viral replication at the cellular level.

The tripartite motif-containing (TRIM) family of proteins plays an important role in restricting viral pathogens and carries out a broad range of immune functions across multiple species.^3,4^ In rhesus macaques, the protein TRIM5, alpha isoform (rhTRIM5*α*) is a potent restriction factor for human immunodeficiency virus (HIV) type 1 (note, HIV refers to HIV type 1 in this work). rhTRIM5*α* interacts with the HIV capsid to strongly restrict HIV, leading to immunity.^5^ The human version of TRIM5*α* (huTRIM5*α*) does not render humans immune to HIV^6^ but does prevent infection by other retroviruses, including N-tropic murine leukemia virus^7,8^ and equine infectious anemia virus.^9^ Human and rhesus TRIM5*α* differ in their ability to restrict HIV despite having high sequence similarity (87% identity, 95% similarity across the whole protein). Introducing mutations in the huTRIM5*α* protein, including a rhTRIM5*α*-like R332P mutation (huTRIM5*α*^R332P^), enables the protein to restrict HIV, but the molecular basis of how these mutations lead to restriction is currently unknown.^10–12^

TRIM5*α* contains four domains: RING, B-box-2, coiled-coil, and PRYSPRY (Figure 1a, Figure S1a). The PRYSPRY domain (Figure 1a,b) is responsible for viral recognition.^13^ It contains thirteen *β*-strands arranged in a *β*-sandwich fold.^14^ Four variable regions (V1-V4) interact with and recognize retroviral capsids, forming the capsid binding surface on one face of the PRYSPRY domain.^14–19^ These regions are rapidly evolving,^20^ exhibiting high sequence variation between species, and determine the specificity of interactions with viral capsids.^19^ Binding partner specificity is highly sensitive to sequence changes in the TRIM5*α* variable regions, as well as in the viral binding partner sequences.^11,14,18,19,21^ The four variable regions either contain or are primarily constituted by loops between *β*-strands, referred to as the V1-V4 loops (Figure S1b and S2, defined here structurally as V1: 326-349, V2: 381-398, V3: 419-428, V4: 483-489, in rhTRIM5*α* residue numbering).

**Figure 1:**
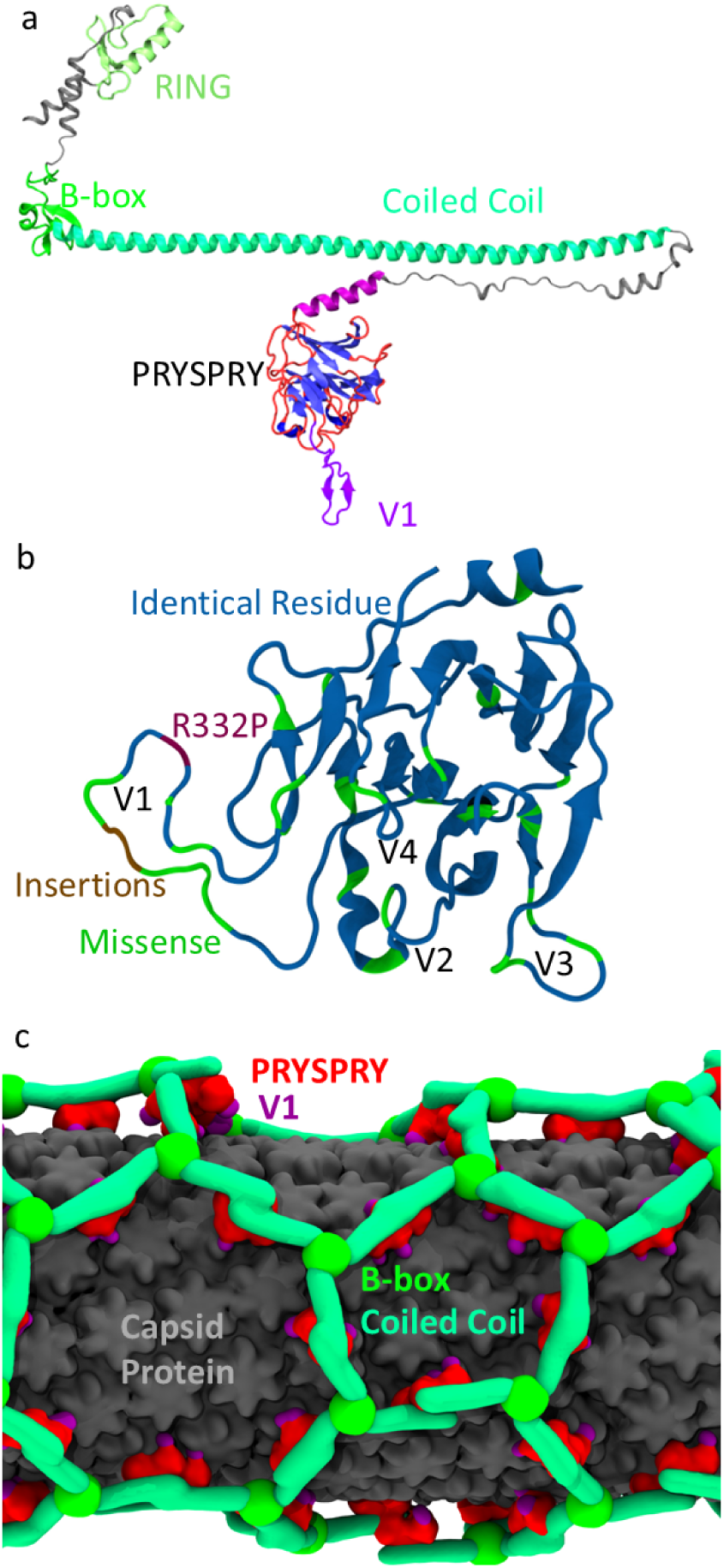
TRIM5*α*, the PRYSPRY domain, and the capsid protein. (a) Structure of the full length huTRIM5*α* protein predicted by AlphaFold Monomer V2.0.^22^ The PRYSPRY domain is colored based on secondary structure (*β*-sheet in blue, *α*-helix in purple, coil in red, and the V1 loop in light purple). The domain boundaries for the RING, B-box and coiled-coil domains defined in Tenthorey *et al.* are used.^10^ These domains are colored in shades of green. Regions not belonging to any domain are shown in gray. (b) The PRYSPRY domain of the rhesus macaque is shown. The structure is taken from our simulations and is colored based on a comparison of the sequences of human and rhesus TRIM5*α*: identical residues in blue, missense mutations in green, inserted residues in brown, and residue R332P (numbered according to huTRIM5*α*) in plum. (c) Structure of a lattice composed of copies of the rhTRIM5*α* protein around a cylindrical tubule composed of copies of the HIV capsid protein. The structure shown is the coarse-grained model based on cryo-electron tomography data from Yu *et al.*^23^ It is colored as follows: capsid protein in gray, PRYSPRY domain in red, V1 loop in purple, coiled-coil domain in pastel green, and B-box domain in lime green.

Early investigations into the relationship between the sequence of TRIM5*α* and HIV restriction identified the N-terminal region of the V1 loop^17^ and another patch within the V1 loop (residues 330-340, in huTRIM5*α* residue numbering)^20^ as being of particular importance. A single point mutation (R332P) allows huTRIM5*α* to restrict HIV almost as potently as rhTRIM5*α*, which also has a proline in the corresponding site on the V1 loop.^11,17^ The deletion of human TRIM5*α* residue R332^12^ and a double mutant (R332G/R335G)^24^ both result in constructs that can restrict HIV more effectively than wild-type, emphasizing the importance of position 332 in the V1 loop. A detailed investigation into the mutational landscape of TRIM5*α* revealed that the majority of random point mutations within the V1 loop of huTRIM5*α* improve its ability to restrict HIV.^10^ The mutations that enhance restriction the most involve removing a positive charge from the V1 loop or adding a negative charge proximal to residue 332 or 335.^10,12^ In fact, mutations that decrease the net charge of the V1 loop were shown to improve HIV restriction in every case.^10^ Reduction in the net charge of the V1 loop, therefore, partially explains the improvement in HIV restriction due to mutations. However, not all of the improvement can be attributed to reduced net charge, as some mutations do not change the charge of the V1 loop but nevertheless lead to improved HIV restriction.^10^ These studies emphasize that HIV restriction potency depends on the sequence of the V1 loop, but the structural basis of this dependency is poorly understood.

Structural information from X-ray crystallography, cryo-electron tomography, and nuclear magnetic resonance (NMR) spectroscopy is available for the HIV capsid protein^23,25–28^ and the rhTRIM5*α* PRYSPRY domain.^14,18^ However, the interaction interface between these proteins involves multiple flexible regions that have been challenging to study experimentally. The surface of the PRYSPRY domain that binds the HIV capsid includes the V1, V2, and V3 loops. The V1 loop has eluded crystallization^18^ and is disordered in both human and rhesus TRIM5*α*. Mutations in the V3 and V4 regions of the PRYSPRY domain can reduce HIV restriction by rhTRIM5*α*.^14^ The HIV capsid is composed of oligomers of the capsid protein (hexamers or pentamers) arranged in a fullerene cone structure.^25^ The capsid protein contains the cyclophilin A (CYPA) binding loop, which undergoes chemical shift changes upon binding the PRYSPRY domain.^27^ Mutations in the CYPA-binding loop allow HIV to resist restriction by both rhTRIM5*α* and multiple mutated variations of huTRIM5*α*.^12,29–31^ Sequence variations in the CYPA-binding loop between subtypes of HIV strongly influence HIV restriction by TRIM5*α*.^32^ Together, these studies provide strong evidence that TRIM5*α-*capsid interactions are dominated by interactions between the CYPA-binding loop and the V1-V4 regions of the PRYSPRY domain.

Molecular dynamics (MD) simulations have been used to investigate the interaction between TRIM5*α* and the HIV capsid protein.^23,33^ Coarse-grained MD simulations showed the spontaneous assembly of a rhTRIM5*α* lattice around the HIV capsid with the PRYSPRY domain interacting mainly with the CYPA-binding loop (refer to Figure 1c for the coarsegrained model of Yu et al.)^23^ However, the nature of coarse-grained simulations precludes the ability to describe the interactions between the proteins in atomic detail. Another study used all-atom MD simulations to obtain a conformational ensemble of the rhTRIM5*α* PRYSPRY domain, followed by docking to predict interactions between highly populated states of the V1 loop and the capsid.^33^ They found that only a subset of V1 conformations led to plausible binding poses with the capsid, and they predicted interactions between the V1-V3 regions of the PRYSPRY domain with the CYPA-binding loop of the capsid protein.^33^ However, docking cannot provide information about the dynamics of the PRYSPRY-capsid interaction interface. Docking yields a static picture, yet such flexible regions are likely to adopt many poses in their interaction. The mounting evidence from both experimental and computational studies indicates that the V1-V4 regions of the PRYSPRY domain and the CYPA-binding loop of the HIV capsid are crucial parts of the interaction interface. An atomistic and dynamic picture of these interactions is still to be determined and is critical for understanding how sequence differences between rhTRIM5*α* and huTRIM5*α* lead to their functional differences in HIV restriction.

Here, we aim to understand the relationship between protein sequence and HIV restriction on a structural level. To this end, we carried out MD simulations of the PRYSPRY domain of rhTRIM5*α* (Figure 2a), huTRIM5*α*, and huTRIM5*α*^R332P^, with a total of over 200 *µ*s of sampling for all systems studied (Table 1). Our simulations reveal that the variable regions of the PRYSPRY domain, especially the V3 loop, are significantly more flexible than previously thought. Through a detailed comparison of huTRIM5*α* and huTRIM5*α*^R332P^ conformational ensembles, we find changes in the local structure near the R332P mutation site. We constructed an atomistic model of the PRYSPRY–capsid complex based on our rhTRIM5*α* simulation ensemble and a coarse-grained cryo-electron tomography model from Yu *et al.*^23^ We performed MD simulations of the complex to provide a dynamic and atomistic picture of the interaction interface between the HIV capsid and the rhTRIM5*α* PRYSPRY domain. Finally, this model for the rhTRIM5*α* and HIV capsid interaction leads to a plausible explanation for the observed differences of rhTRIM5*α*, huTRIM5*α*, huTRIM5*α*^R332P^ in HIV restriction. Our model indicates that repulsive interactions between residue R82 of the capsid protein and positively charged residues on the huTRIM5*α* V1 loop prevent capsid binding and, therefore, restriction of HIV by huTRIM5*α*. In support of this model, we show that mutating these positively charged residues to a neutral residue (either TRIM5*α*-R332 or capsid-R82) reduces this repulsion, thereby enhancing binding to the HIV capsid and restriction by huTRIM5*α*^R332P^.

**Figure 2:**
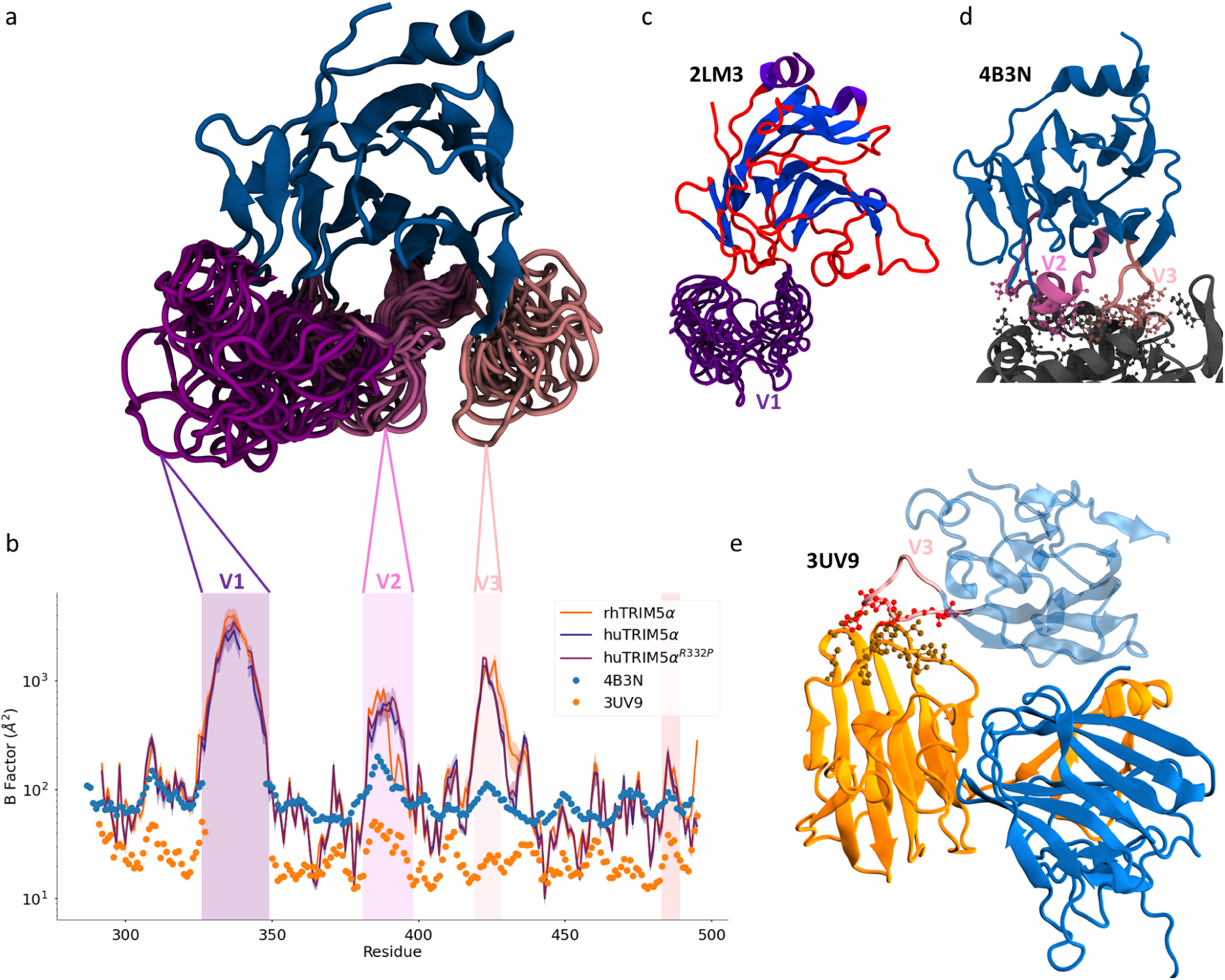
The PRYSPRY domain contains three flexible loops. (a) A 3D structure of the rhTRIM5*α* PRYSPRY domain taken from simulations. Multiple conformations of the V1-V3 loops are shown superimposed. The structure is shown in cartoon representation. The V1 loop is shown in dark purple in panels a, b and c, the V2 loop is shown in mauve in panels a and d, and the V3 loop is shown in pink in panels a, d, and e. (b) B-factors for rhTRIM5*α*, huTRIM5*α*, and huTRIM5*α*^R332P^ PRYSPRY domain as computed from simulation trajectories (see Equation 1). Shading indicates the standard error of the mean obtained by treating each trajectory as independent. B-factors of C*α*-atoms from the two available crystal structures (PDB IDs 4B3N,^18^ 3UV9^14^) are also shown. Values are plotted on a logarithmic scale. The vertical shaded areas colored in purple, mauve, pink, and red indicate the locations of the V1, V2, V3, and V4 loops, respectively. (c) NMR ensemble (PDB ID: 2LM3^14^) of the PRYSPRY domain of rhTRIM5*α*. The protein is colored by secondary structure (*β*-sheet, blue; *α*-helix, purple; coil, red) and the V1 loop structures are shown in dark purple. (d) Crystal structure of the PRYSPRY domain of rhTRIM5*α* (PDB ID: 4B3N^18^). Residues forming crystal contacts involving the V2 and V3 loops are shown in ball-and-stick representation. (e) Crystal structure of the PRYSPRY domain of rhTRIM5*α* with the V1 loop removed (PDB: 3UV9^14^), which crystallized as a domain-swapped trimer. Individual PRYSPRY domains are colored differently. The V3 loop of one of the PRYSPRY domains is shown in pink, with crystal contacts formed by the V3 loop shown in ball and stick representation.

**Table 1:**
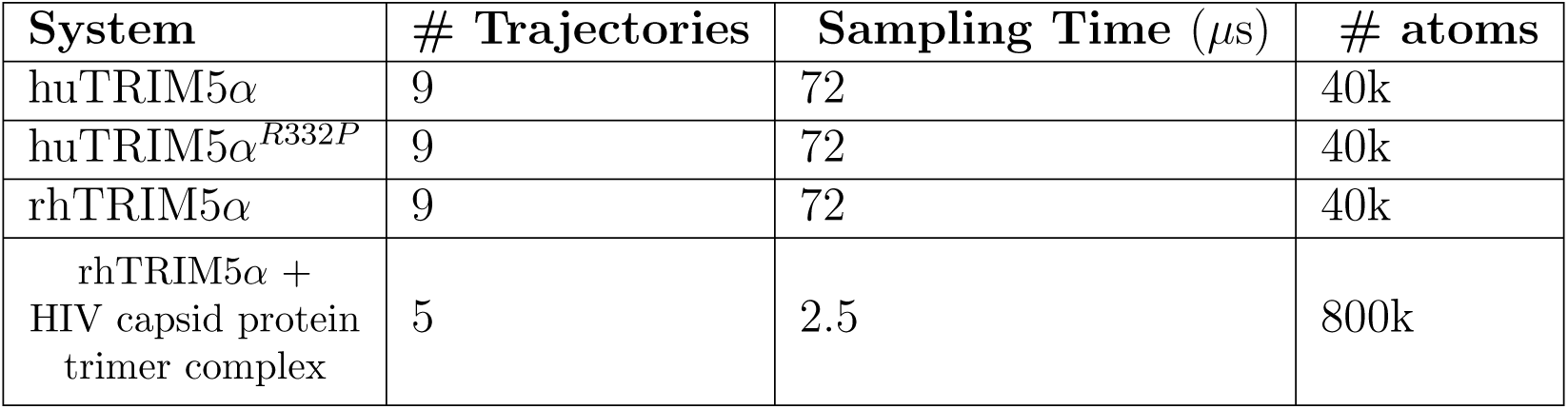
List of simulations performed in this study.

## Results

### V1, V2, and V3 loops exhibit high structural flexibility

We sought to uncover the molecular basis for how TRIM5*α* evolves improved antiviral function against retroviruses, including HIV. To do so, we conducted all-atom simulations of the virus-binding PRYSPRY domain of TRIM5*α* variants with differing degrees of function against HIV (see Methods for details). The four variable regions of TRIM5*α* exhibit sequence diversity across species and form the binding interface with the capsids of retroviruses, including HIV.^16,18,34,35^ Within each of these variable regions are loops, which we refer to as the V1-V4 loops (refer to Figure S1b for the sequences of the variable loops and regions). In both human and rhesus TRIM5*α*, the V1 loop is the longest (22 residues in huTRIM5*α* and 24 residues in rhTRIM5*α*). The V2 loop has 18 residues, and the V3 loop has 10 residues in both species.

The V1 loop of the PRYSPRY domain of rhTRIM5*α* is known to be disordered.^14^ We hypothesized that the degree of V1 flexibility might explain differences in restriction. To quantify the flexibility of the V1 loop in solution across TRIM5 orthologs, we computed the root-mean-square fluctuation (RMSF) from each of our simulation ensembles. From the RMSF profiles, we computed the corresponding B-factors, a metric of per-residue positional variability (using Equation 1, see Methods). These B-factors are shown alongside the B-factors reported in the two available crystal structures of rhTRIM5*α*^14,18^ in Figure 2b. Consistent with a previous NMR study that reported V1 loop disorder^14^ (PDB ID: 2LM3,^14^ Figure 2c), the B-factors of the simulation ensemble of rhTRIM5*α* are highest in the V1 loop. In the crystal structures of rhTRIM5*α*, the V1 loop is either not observed (PDB ID: 4B3N,^18^ Figure 2d) or is replaced by a two-residue linker to enable crystallization (PDB ID: 3UV9,^14^ Figure 2e), consistent with its flexibility.

We also observe high B-factors in the V2 and V3 loops across all TRIM5 variants, indicating that all three of these loops are flexible (Figure 2a-b). The two crystal structures of rhesus TRIM5 show some differences in the conformation of the V2 and V3 loops (Figure 2d-e, Figure S3), suggestive of some degree of conformational heterogeneity. However, the B-factors computed from our simulated ensembles are much higher than the crystallographic B-factors for residues in the V2 and V3 loops (Figure 2b). It is, of course, to be expected that the protein will exhibit a higher flexibility in solution compared to a crystal environment.^36^ Differences between the simulated and experimental B-factors in the V2 and V3 loops can be explained by the fact that both of these loops form crystal contacts with neighbouring proteins in the crystal^14,18^ (Figure 2d-e). Residues in the V4 loop exhibit much lower flexibility compared to the V1-V3 loops (Figure 2b).

In addition to crystallographic B-factors, we reviewed other experimental data that could provide further insight into the flexibility of the V1-V3 loops. Previously published ^1^H-^15^N nuclear Overhauser effect (NOE) data are consistent with high mobility of the V1 and V3 loops in rhTRIM5*α*.^14^ NOEs were not observed for several residues in the V2 loop, which could indicate high flexibility in this loop.^14^ Our simulated ensemble also correlates well with observed chemical shifts^14,37^ for rhTRIM5*α* across all residues, including within the variable loops (Figure S4). These NMR data support our observation that the V1-V3 loops are flexible.

Because the V1-V3 loops exhibit such high flexibility in our simulations, we asked if these regions would be predicted to be disordered based on their sequence composition. We obtained a prediction for the propensity of each residue to be disordered in TRIM5*α* (Figure S5) using Metapredict Version 2.^38,39^ Unexpectedly, the V1, V2 and V3 loops from both species’ TRIM5 are predicted to be ordered by Metapredict. Structure predictions made by AlphaFold include predicted local difference distance test (pLDDT) scores, which describe the confidence in the predicted structure. pLDDT scores can also be used to predict disordered regions.^40,41^ To compare with the disorder prediction of Metapredict, we computed a normalized disorder prediction score from the AlphaFold pLDDT scores (1-pLDDT/100) (Figure S5). Using this normalized pLDDT score as a disorder prediction, we find that the V1-V3 loops are all above baseline. In fact, the normalized pLDDT scores are remarkably consistent with the flexibility in these regions observed in simulations (shaded regions in Figure 2b, Figure S5). A previous simulation study also found that the V3 loop is flexible, but considerably less flexible than the V1 loop.^33^ The fact that we observe comparable flexibility between the V1 and V3 loops could be explained by the more extensive sampling in the current study.

The flexibility of the V1 loop has led to speculation that loop flexibility may permit recognition of distinct capsid proteins through variation in the loop’s conformations.^33^ Since proline tends to promote disorder,^42^ one hypothesis is that the R332P mutation in huTRIM5*α* may promote disorder in the V1 loop and improve recognition of the HIV capsid protein via enhanced flexibility. However, we observe minimal differences in the flexibility across the protein for all three systems (Figure 2b). Some minor differences exist between the rhesus and human systems. Nevertheless, huTRIM5*α* and huTRIM5*α*^R332P^ have nearly identical flexibility across the entire PRYSPRY domain. We conclude that there is no evidence from our simulations for increased disorder due to the R332P mutation contributing to improved HIV restriction in huTRIM5*α*^R332P^.

### Conformational changes in huTRIM5*α* due to the R332P mutation

Mutation of residue R332 to proline is among the most effective point mutations in improving huTRIM5*α*’s ability to restrict HIV.^10^ Tenthorey *et al.* found that positive charge in the V1 loop is the dominant impediment to HIV restriction.^10^ However, it is not clear whether the removal of positive charge improves HIV restriction by affecting electrostatic interactions with the HIV capsid or through a charge-dependent conformational change within the PRYSPRY domain itself. Furthermore, the removal of positive charge cannot explain all of the experimentally observed improvement in HIV restriction, since there is variation in restriction even between mutations that do not reduce the net charge. For example, mutation of G333 to almost any other residue, including those that do not affect the net charge, improves HIV restriction considerably.^10^ Mutation-induced conformational changes within the PRYSPRY domain could explain these charge-independent differences.

To investigate the possibility that conformational changes in the PRYSPRY domain cause the observed improvements in HIV restriction, we compared the conformational ensembles of the huTRIM5*α* and huTRIM5*α*^R332P^ simulations. We first analyzed the conformational ensembles with respect to the prevalence of different types of turns. Proline residues are statistically common in *β*- and *γ*-turns.^43,44^ The structures of huTRIM5*α* and rhTRIM5*α* predicted by AlphaFold^22^ both include a *β*-hairpin in the V1 loop (Figure 3a,b). At the corner of this hairpin is residue R332 in huTRIM5*α*, which corresponds to residue P334 in rhTRIM5*α*. In the sequences of both huTRIM5*α* and rhTRIM5*α*, the following residue is a glycine. Because the sequence motif PG is known to promote *β*-turn formation,^45,46^ we investigated whether the R332P mutation affects the probability of turn formation in huTRIM5*α*. To this end, we computed the distance between the C*α* atoms on either side of the mutation site in our simulation ensembles (Figure 3a-c). The distributions of these distances are bimodal, indicative of a closed and open state. Both huTRIM5*α*^R332P^ and rhTRIM5*α* have an increased population of the closed state compared to huTRIM5*α*, indicating that a proline in position 332 promotes a turn conformation over an extended conformation.

**Figure 3:**
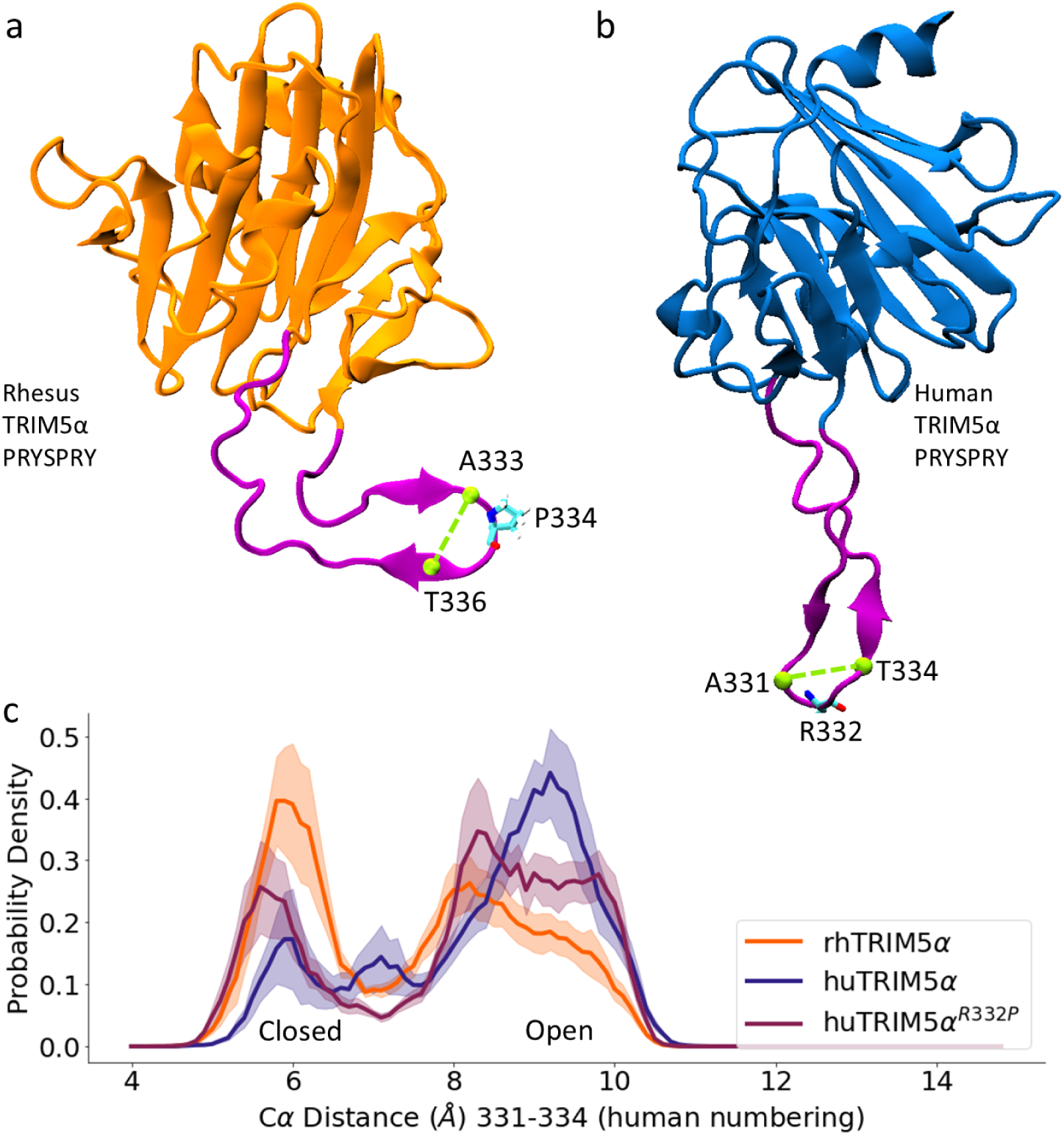
*β*-hairpin predicted by AlphaFold and stabilized by the R332P mutation. (a-b) The structure of rhTRIM5*α* (a) and huTRIM5*α* (b) PRYSPRY domain predicted by AlphaFold V2.0^22^ are shown in cartoon representation. The V1 loops are shown in purple. huTRIM5*α* residue R332 and the corresponding residue P334 in rhTRIM5*α* are shown in licorice representation. In (a), residues A333 and T336 C*α* atoms are shown as green spheres. In (b), residues A331 and T334 C*α* atoms are shown as green spheres. In (a) and (b), the structures are in different relative orientations to highlight the V1 loop, which points in different directions in each structure. (c) Probability distribution of the distance between the C*α* atoms of residues A331 and T334 (A333 and T336 in rhTRIM5*α*) from the simulations. Shading indicates standard error of the mean obtained by treating each trajectory as independent.

Structural differences due to the proline at this position can also be seen in the distance difference map and contact difference map of the V1 loop (Figure S6a,b). The contact map indicates that contacts between residues 331 and 333-335 are more probable in TRIM5*α*^R332P^ compared to wild type. In a contact map between residue pairs, a *β*-hairpin structure corresponds to contacts perpendicular to the diagonal. Such a structure is faintly visible in the contact maps of the V1 loop in all three systems (Figure S6c-e), suggesting that although the *β*-turn is more prevalent in huTRIM5*α*^R332P^ compared to huTRIM5*α*, a *β*-hairpin-like structure is present with low probability in all three systems. Furthermore, these results demonstrate that the occupancy of the *β*-turn generally correlates with antiviral function.

We next asked whether other structural differences in the PRYSPRY domain might explain the effect of the human TRIM5*α* R332P mutation on antiviral function. We computed various structural properties, including secondary structure (Figure S2), radius of gyration (R*_g_*) (Figure S7), and contacts (Figure S8). We observe very little difference in the secondary structure of the PRYSPRY domain (Figures S2), except for a slight increase in the *β*-propensity of two residues in the V1 loop for R332P relative to wild type (Figure S2c). There is an increased helix propensity observed in the V2 loop in rhTRIM5*α* compared to huTRIM5*α* (Figure S2a). Although these differences are statistically significant, there is no clear relationship between them and any improved HIV restriction. The R*_g_* distribution of the V1 loop is polymodal, but with large uncertainty (Figure S7). We observe a very slight, but statistically significant, difference in the V1 loop R*_g_* distribution between huTRIM5*α* and huTRIM5*α*^R332P^. We also observe a polymodal distribution with large uncertainty in the R*_g_* of the V3 loop. The mean R*_g_* is the same within uncertainty, but the distributions of R*_g_* differ. The V2 loop has a slightly larger R*_g_* in huTRIM5*α*^R332P^. Overall, both the R*_g_* distributions and the contact difference map (Figure S8) show relatively small differences in structure across the entire protein due to the R332P mutation, making it unlikely that these changes could explain the dramatically improved restriction of HIV.

The conformational differences we observe between the PRYSPRY domain of huTRIM5*α* and huTRIM5*α*^R332P^ are largely minimal, with the exception of the stabilization of the turn and *β*-hairpin-like structure near the location of the R332P mutation. We hypothesize that the increased turn propensity at the tip of the loop induced by the R332P mutation could promote insertion of the V1 loop into the space between HIV capsid hexamers. It could also be that the formation of the turn leads to specific interactions with the capsid that are only possible with this conformation. While we cannot rule out these conformational differences as contributors, it seems likely that they are not the most important factors contributing to the improved HIV restriction by TRIM5*α*^R332P^. This view is consistent with the finding that the largest impediment to HIV restriction is the removal of positive charge,^10^ which suggests that electrostatic interactions between the HIV capsid and the PRYSPRY domain play a critical role. We explore this latter hypothesis and find good evidence to support it below.

### Clustering of the rhTRIM5*α* Loop States

To understand how the TRIM5*α* PRYSPRY domain interacts with the HIV capsid, we focused our subsequent efforts on the most potent antiviral variant, rhTRIM5*α*. As a first step, we sought to obtain representative conformations of the PRYSPRY domain of the rhTRIM5*α* ensemble to use as starting points for our simulations. To this end, we constructed a conformational landscape of the rhTRIM5*α* PRYSPRY domain using principal component analysis (PCA). Since most of the protein is well-structured, we focused on the flexible V1V3 loops to characterize the conformational ensemble. We also chose to focus on these loops because, as discussed above, there is substantial evidence from prior studies that they interact directly with the capsid. We performed PCA on the pairwise distances of the C*α* atoms of a selection of residues encompassing the flexible V1-V3 loops to create a two-dimensional conformational landscape. The resulting conformational landscape of the PRYSPRY domain of rhTRIM5*α* is shown in Figure 4a and Figure S9. We used density-based spatial clustering of applications with noise (DBSCAN)^47^ to cluster the most densely populated conformational states in this landscape (Figure 4a).

**Figure 4:**
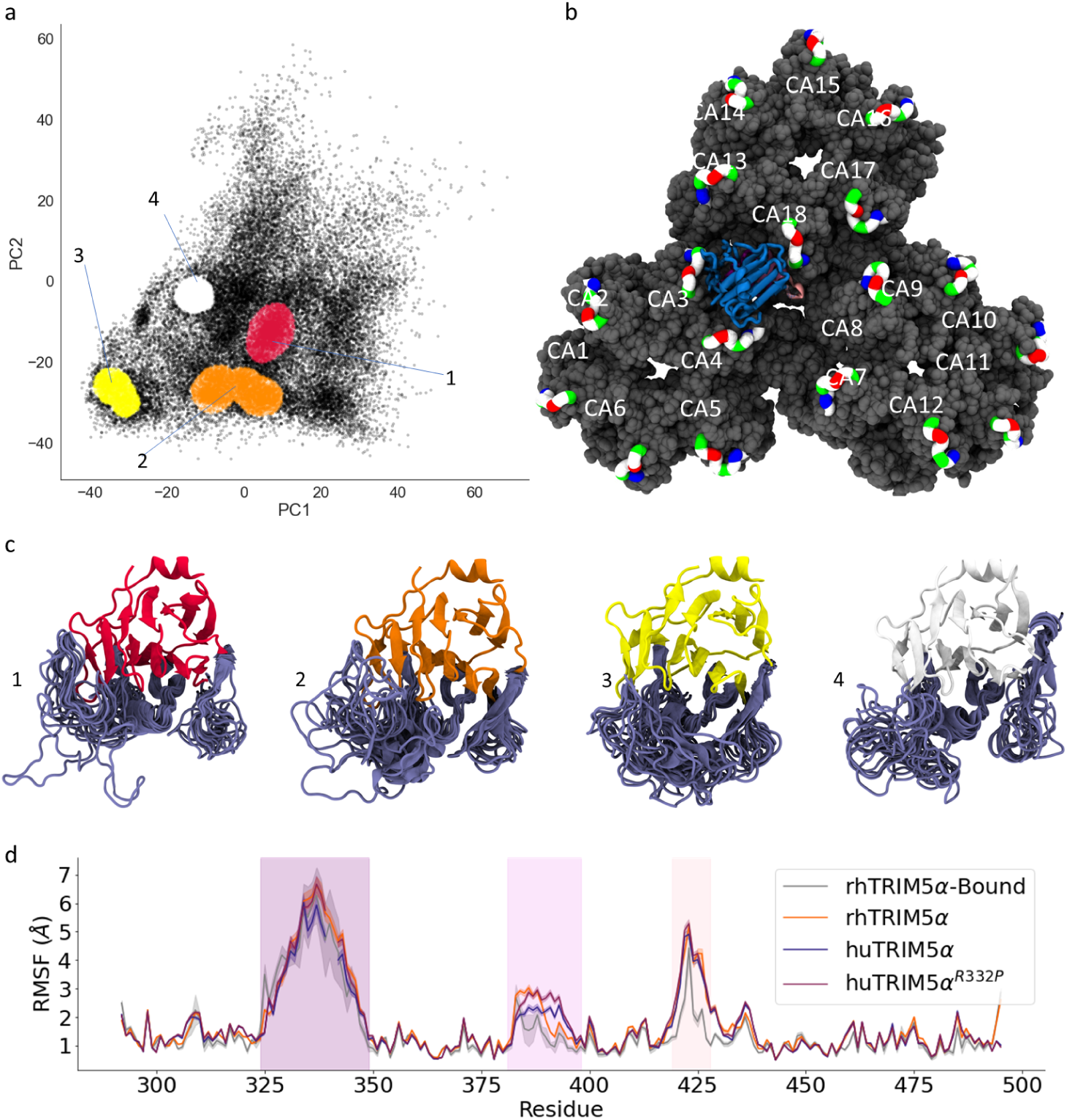
Flexibility and clusters of the rhTRIM5*α* loops. (a) PCA of the rhTRIM5*α* PRYSPRY domain ensemble with DBSCAN clustering of the conformational landscape. Each point in the 2D space defined by the first two principal components (PC1 and PC2) corresponds to a structure from the rhTRIM5*α* PRYSPRY domain simulations (see Methods for description of the PCA). The four clusters are shown in red (1), orange (2), yellow (3), and white (4), and the outliers are shown in black. (b) 3D structure of rhTRIM5*α* PRYSPRY domain (blue, with the V1, V2, and V3 loops shown in purple, mauve, and pink, respectively) in complex with a trimer of hexamers of the capsid protein (CA, shown in gray). The 18 individual capsid monomers are labelled with assigned numbers. The CYPA-binding loop of each capsid monomer is emphasized and colored by residue type (white: hydrophobic, green: polar, red: negatively-charged, blue: positively-charged). (c) Structural ensembles of the four loop state clusters in (a). V1-V3 loops are colored in violet. 15 structures selected at random from each cluster are shown. (d) RMSF profiles of all TRIM5*α* simulation ensembles. The RMSF of the HIV capsid-bound rhTRIM5*α* PRYSPRY domain is shown in gray together with the RMSF of the unbound systems. To compare fairly with rhTRIM5*α* bound to the capsid, an equal amount of simulation time was used to compute the RMSF profile of the unbound systems. That is, the RMSF profile was computed over 500 ns intervals and averaged. Shaded areas above and below the lines indicate the standard error of the mean treating each 500 ns interval as an independent measurement. Vertical shaded regions indicate the locations of the V1 V2, and V3 loops.

The clustering analysis resulted in four densely populated states. Representative structural ensembles from each of these states are shown in Figure 4c. The four clusters differ in terms of both the relative loop positioning and the structure of the loops. We next investigated the degree to which structures from these clusters can be docked onto the HIV capsid without steric clashes. We used a multimeric assembly of capsid proteins, a trimer of hexamers (Figure 4b), in place of the entire HIV capsid, which we refer to simply as the HIV capsid henceforth. We chose one structure at random from each cluster and found that only one of these structures (cluster 3, Figure 4a,c) fits onto the HIV capsid without any steric clashes between the proteins (see Figure S10 for an example of steric clashes in the aligned complex). We subsequently used this structure for MD simulations of the complex (Figure 4b). Additional details of the methodology used to obtain an initial structure for the rhTRIM5*α*-capsid complex are provided in Methods. We note that the clustering analysis used here is not unique. Different clusters would be expected using different clustering algorithms or parameter choices. However, the clustering here is sufficient for our purpose, namely, to provide representative conformations of rhTRIM5*α* that can be docked to the HIV capsid.

### The V1-V3 loops remain flexible upon HIV capsid binding

To investigate changes in the PRYSPRY domain of rhTRIM5*α* upon capsid binding, we performed MD simulations of the rhTRIM5*α*-HIV capsid complex (Figure 4b, see Methods). rhTRIM5*α* has weak affinity for the HIV capsid protein, which is thought to be enhanced by avidity effects in which rhTRIM5*α* restricts HIV by forming higher-order oligomers that envelope the HIV capsid.^21^ However, the rhTRIM5*α* PRYSPRY domain has been shown to be sufficient in its monomeric form to recognize and bind to the HIV capsid protein.^18^ For this reason, we simulated the rhTRIM5*α* PRYSPRY domain as a monomer in complex with the HIV capsid. Five independent simulations 500 ns in length were carried out. We note that on this relatively short timescale, the PRYSPRY domain does not dissociate from the capsid, or sample all possible conformational states of the V1-V3 loops. We computed the RMSF of the PRYSPRY domain in the complex and compared it to the RMSF in the unbound simulations (Figure 4d). The V1-V3 loops retain high flexibility in the complex, with all three loops having a higher RMSF than the ordered regions of the PRYSPRY domain. There is a slight decrease in the RMSF of the V2 and V3 loops compared to the unbound simulations. Overall, these results indicate that the loops remain flexible and disordered when interacting with the capsid protein.

Next, we investigated how interactions with the HIV capsid influence the conformational ensemble of the PRYSPRY domain. To do so, we projected the capsid-bound rhTRIM5*α* trajectories onto the conformational landscape of the unbound rhTRIM5*α* used for clustering (Figure S11). The trajectories sample conformations similar to the cluster used as an initial structure, but they also explore some of the other regions of the landscape. The adherence to the initial structure may be due to the limited sampling of the simulations of the complex. We note that the conformations explored by the V1-V3 loops of rhTRIM5*α* when bound to the capsid are largely within the bounds of the conformations sampled in the unbound simulations, suggesting similarity of the bound and unbound ensembles. On this basis, and consistent with Kovalskyy and Ivanov’s model for capsid recognition based on docking,^33^ we suggest that the bound rhTRIM5*α* ensemble represents a subset of the unbound ensemble.

### The CYPA-binding loop and the variable loops in the PRYSPRY domain mediate the capsid-TRIM5*α* binding interface

A detailed description of the interaction interface between the HIV capsid and the TRIM5*α* PRYSPRY domain interface is critical to understand how the sequence differences between rhesus and human TRIM5*α*, as well as mutations in human TRIM5*α*, lead to the differences in HIV restriction. To characterize this interface, we started by computing the contact frequency by residue between the capsid and the TRIM5*α* PRYSPRY domain (Figure 5a, Figure S12, Figure S14). The PRYSPRY domain is situated at the interface between three capsid hexamers (Figure 4b). It interacts with five of the eighteen capsid monomers during the simulations. All of the contacts formed are highly transient, and none persist throughout all simulation trajectories, which is evident from the low values of the contact propensities (Figure S12). Instead, contacts are repeatedly formed and broken between multiple residue pairs. No specific binding interactions are stable, and instead, a large multiplicity of varied interactions hold the complex together. Several conformations of the complex are shown in Figure S13.

**Figure 5:**
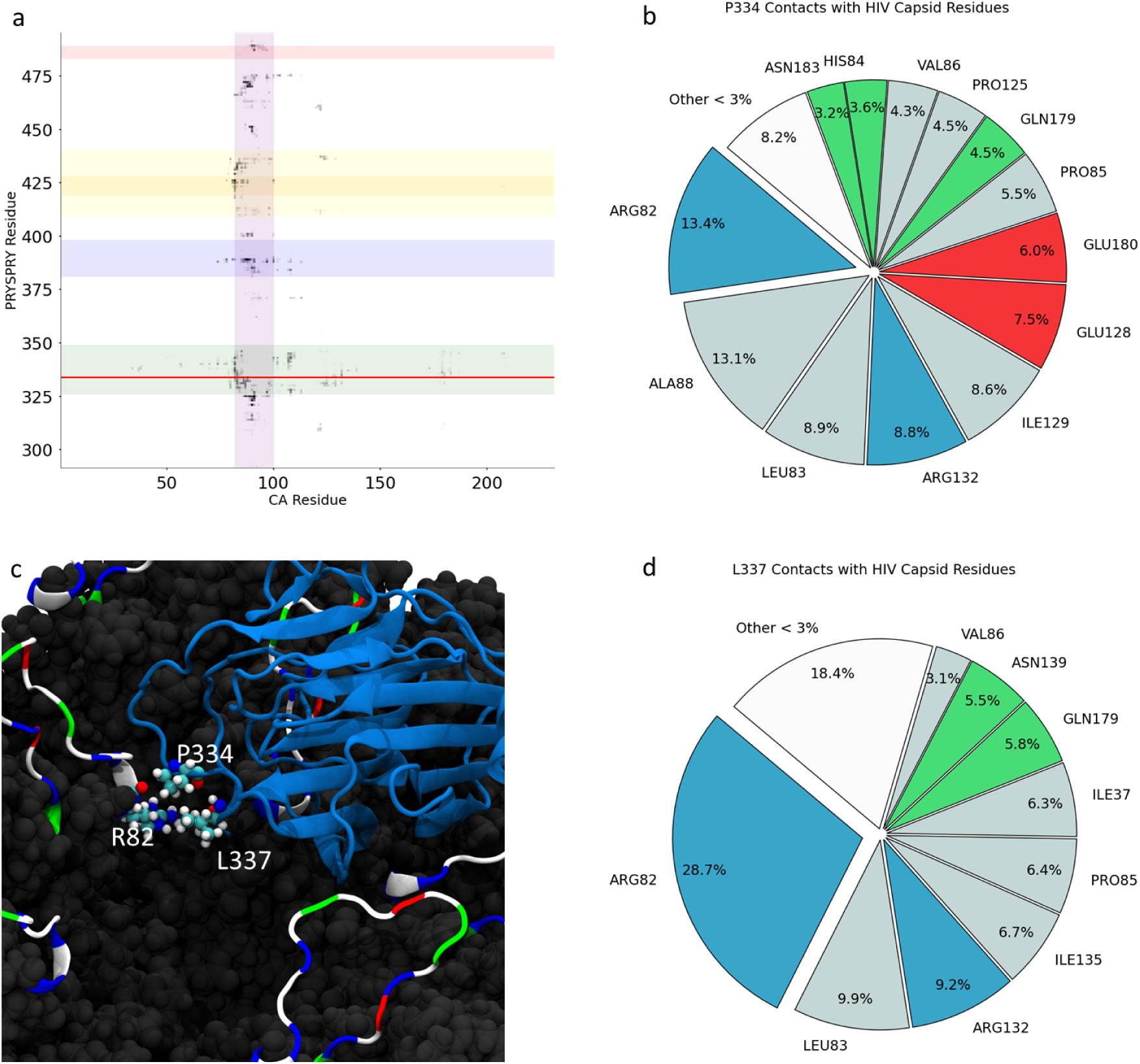
Contacts formed between rhTRIM5*α* and the HIV Capsid. (a) Relative contact propensities between rhTRIM5*α* and HIV capsid protein, averaged over all 18 capsid monomers, are indicated with increasing color for each residue pair. Contact propensities represent the probability for each contact to occur; these values range from 0 to 0.022. Shading delineates various regions of each protein (green: V1 loop (326-349), blue: V2 loop (381-398), orange: V3 loop (419-428), yellow: the broader V3 variable region, red: V4 loop (483-489), purple: the loop of the HIV capsid protein where CYPA binds (capsid protein residues 82-100)). The red horizontal line indicates the location of residue P334 (corresponding to R332 in huTRIM5*α*). (b) All HIV capsid protein residues that residue P334 of rhTRIM5*α* forms contacts with are shown according to their respective portion of all P334-capsid contacts. (c) A structure of the complex from simulations is shown, focusing on the contact formed between L337 and P334 on the PRYSPRY domain and R82 on the CYPA-binding loop of the capsid protein. The PRYSPRY domain is shown in blue and the HIV capsid is shown in dark gray. The CYPA-binding loop of each capsid protein is shown in a cartoon representation colored by residue type (white: hydrophobic, green: polar, red: negatively-charged, blue: positively-charged).(d) All HIV capsid protein residues that residue L337 of rhTRIM5*α* forms contacts with are shown according to their respective portion of all L337-capsid contacts.

We computed the contacts formed between the PRYSPRY domain and HIV capsid, averaging over the 18 capsid monomers present in the simulations of the complex. We find that the vast majority of contacts are formed with the CYPA-binding loop of the capsid protein, and the majority of the contacts formed by the PRYSPRY domain involve residues in the V1-V3 loops (Figure 5a, Figure S14). This characterization of the interaction interface is highly consistent with previous experimental and computational studies, which have implicated the CYPA-binding loop and the variable regions of the PRYSPRY domain as being the dominant participants in the interaction.^10–12,14,17,20,23,24,27,29–31,33^ In particular, Sawyer et al. identified a patch within the V1 loop (residues 330-340, in huTRIM5*α* numbering) harbouring several residues exhibiting positive selection.^20^ This is significant because such dense regions of positive selection are likely to be contact points between co-evolving viral and antiviral proteins.^20^ In our simulations, all residues in the patch identified by Sawyer et al. form contacts with the HIV capsid (Figure S14a), suggesting that this patch is indeed a point of physical contact between the two proteins. The transiency of the interactions we observe, along with the retention of flexibility of the loops upon binding, is indicative of a fuzzy binding interface between the two proteins.

### Gain-of-function mutation sites of TRIM5*α* interact with positively charged residues on the HIV capsid

We next use our simulation of the complex to understand why point mutations in human TRIM5*α* exhibit improved recognition of HIV. We focused our analysis on residues R332 and R335, the positions at which the most potent gain of function mutations in TRIM5*α* are found.^10–12^ These residues correspond to P334 and L337 in rhTRIM5*α*. Both P334 and L337 interact with many capsid protein residues in our simulations of the complex, reflecting the conformational heterogeneity of the entirety of the PRYSPRY-HIV capsid interface (Figure 5b,d). Importantly, P334 and L337 both form contacts most frequently with residue R82 on the CYPA-binding loop. This observation is consistent with NMR experiments that indicated chemical shift changes of residue R82 in cross-linked capsid hexamers upon complex formation with rhTRIM5*α*.^27^ Based on the results of our simulations, and the finding that any charge-neutralizing mutations at or near R332 of human TRIM5*α* improve its restriction of HIV,^10–12^ we hypothesize that the improvement in huTRIM5*α*’s ability to restrict HIV as a consequence of the R332P or R332G/R335G mutations is largely due to electrostatic effects, primarily with R82. Removing the positive charge from TRIM5 at those sites would make interaction with the arginine on the capsid (R82) more favourable. Mutagenesis studies of huTRIM5*α* indicate that removing positive charge or introducing negative charge near these sites in the V1 loop significantly enhances HIV restriction, whereas the R332K mutation does not.^10,12,24^

A prediction based on these results is that neutralizing the positive charge on either human TRIM5*α* or the HIV capsid should increase TRIM5*α*-mediated restriction, as either mutation would be sufficient to remove the electrostatic repulsion, whereas neutralizing both charges simultaneously should not provide any additive benefit. Indeed, HIV has reduced infectivity in human cells when the capsid acquires a mutation of R82L,^30,48^ which might be explained by huTRIM5-mediated restriction. To test this prediction directly, we made several mutations at position R82 of the HIV capsid that either neutralized (leucine, L; methionine, M) or retained (lysine, K) the positive charge. We then measured the susceptibility of these HIV variants to huTRIM5*α*-mediated restriction. Consistent with our prediction, mutations that neutralized the charge increased wild-type human TRIM5*α*’s ability to restrict HIV, whereas a charge-conserving R82K mutation did not (Figure 6). Importantly, mutations at R82 did not significantly affect restriction by the R332P variant of human TRIM5*α*, indicating that their effect disappears once electrostatic repulsion is removed through alternative means. In contrast, huTRIM5*α* with the R332P mutation has an additional ∼2 fold improved restriction of all HIV capsid variants tested when compared to wild-type TRIM5 restriction of R82L or R82M, which might be explained by the improved turn propensity of this variant (Figure 3c). Notably, our simulations of the complex indicate low prevalence of the closed turn conformation (Figure S6f) compared to the apo rhTRIM5*α* simulations. However, the initial state of the complex simulations is in an open state, which could bias the overall distribution given the limited conformational sampling. Consistent with the importance of this electrostatic repulsion for HIV escape from human TRIM5*α*, the R82 position is 99.5% conserved among HIV-1 isolates, and the only repeatedly observed variant at this site is a positively charged lysine.^49^ In contrast, another strain of HIV, HIV type 2, has a valine residue in place of R82^50^ and has also been shown to be susceptible to huTRIM5*α*.^51^ These experimental observations, combined with our simulation data, are consistent with charge repulsion between R82 and R332/R335 being the dominant impediment to HIV restriction by huTRIM5*α*.

**Figure 6:**
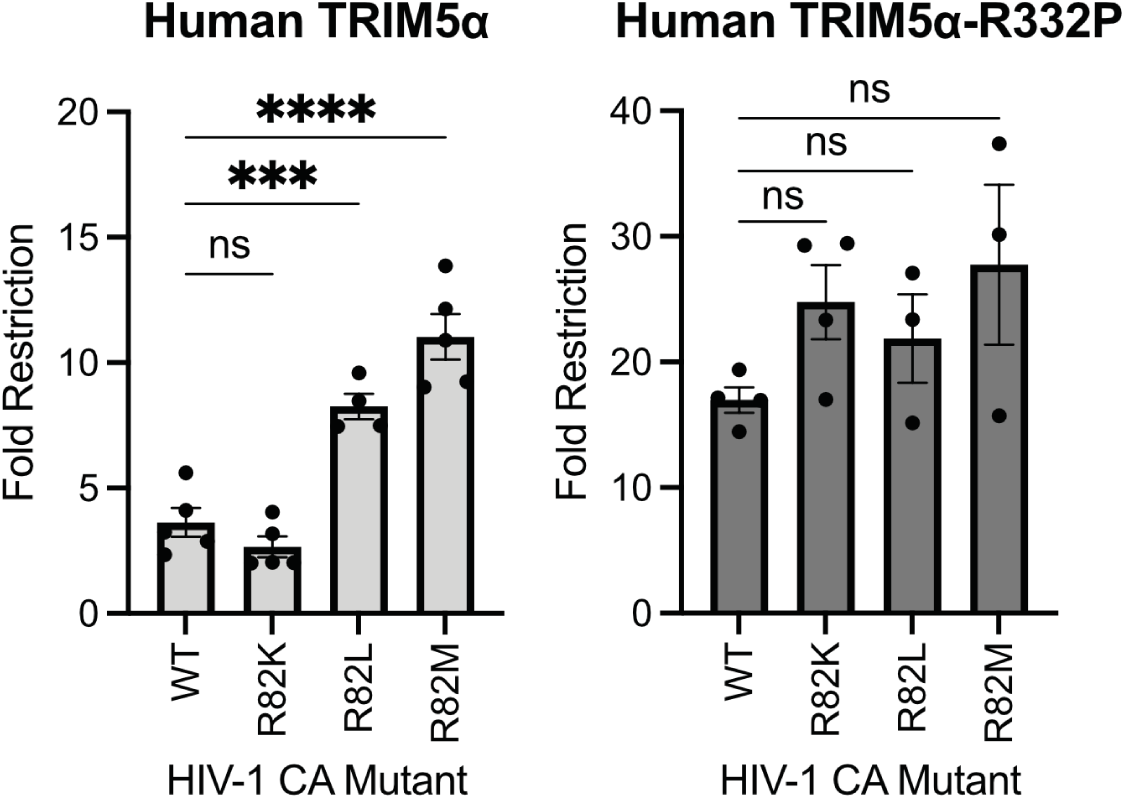
huTRIM5*α* restriction improves when R82 is mutated to a non-polar residue. HIV capsid mutants were used to infect CRFK cells expressing either wild-type or an R332P mutant of human TRIM5. TRIM5-mediated fold restriction was determined by measuring the decrease in infection relative to CRFK cells expressing a matched empty vector (see Methods). Data are representative of at least 3 independent experiments. Error bars indicate the standard error of the mean. ns indicates not significant; *** indicates p *<* 0.001; and **** indicates p *<* 0.0001 using one-way analysis of variance with Dunnett’s multiple comparisons correction vs. wild-type HIV.

Somewhat unexpectedly, the residue in the rhesus PRYSPRY domain that forms contacts with the HIV capsid most frequently is Y389, which resides not in the V1 loop but in the V2 loop (Figure S14a). Several other residues outside of the V1 loop also form frequent contacts with the capsid (Figure S14a). The most prominent contacts Y389 forms are with residues in the CYPA-binding loop (A88, H87, G89, I91, Figure S15). Residues G89 and I91 have been shown to undergo chemical shift changes upon rhTRIM5*α* binding.^27^ Y389 is also one of two residues in the V2 loop that differs between rhesus and human TRIM5*α* (it corresponds to C385 in huTRIM5*α*, Figure S1b). Since any sequence difference between the two proteins could potentially contribute to differences in HIV restriction, this difference also hints at the importance of residue Y389 in viral capsid recognition. Furthermore, a mutation study of orangutan TRIM5*α* indicated that this residue (corresponding to Y385 in orangutan TRIM5*α*) is involved in determining viral restriction specificity.^19^ We speculate that sequence differences between human and rhesus TRIM5*α* outside the V1 loop could also contribute to the observed differences in HIV restriction.

## Discussion

The present study provides a probabilistic description of the ensembles of multiple variants of TRIM5*α* towards a structural understanding of the functional differences between them. Our simulations reveal that the V2 and V3 loops are more flexible than has been previously observed. They also indicate changes in the conformational ensemble of the V1 loop of huTRIM5*α* due to the R332P mutation. Specifically, we observe an increase in the probability of a *β*-turn including residue 332 in huTRIM5*α*^R332P^ and rhTRIM5*α* compared to huTRIM5*α*. We hypothesize that these structural differences could be relevant to the improved HIV restriction observed in huTRIM5*α*^R332P^ compared to other mutations of residue R332, and rhTRIM5*α* compared to huTRIM5*α*. Evaluating the potential involvement of the *β*-turn and the associated *β*-hairpin-like structure in capsid binding will require further structural and mutation studies.

We also carried out the first atomistic MD simulations of the TRIM5*α*-HIV capsid complex, revealing information about the fuzzy interaction interface of the two proteins. Consistent with previous studies,^10–12,14,17,20,23,24,27,29–31,33^ we demonstrate that the interaction interface primarily consists of the CYPA-binding loop and the variable regions of the PRYSPRY domain. We also provide a plausible explanation for the improved HIV restriction observed in multiple mutants (including R332P, R332G/R335G) of the huTRIM5*α* protein. This mechanism, which involves more favourable electrostatic interactions with residue R82 on the HIV capsid CYPA-binding loop, is highly consistent with previous experimental data demonstrating the importance of positive charges in the V1 loop of huTRIM5*α* in hindering HIV restriction.^10,12,24^ It has previously been speculated that electrostatic repulsion between the CYPA-binding loop and positive residues in the V1 loop impair huTRIM5*α* binding to the HIV capsid,^10^ which is consistent with the mechanism suggested by our simulation results and our experimental data. Finally, we find that the loops remain flexible while interacting with the capsid, indicating that there are likely multiple, heterogeneous interaction modes relevant to TRIM5*α*-capsid binding.

While our study provides a first view of the HIV capsid-TRIM5*α* interface in all-atom detail, it does so with limited sampling of the possible conformational space of the system. Given that the variable loops remain flexible upon binding the capsid protein, we emphasize that the contact probabilities computed from our simulations may not reflect the true distributions due to limited sampling. The sampling problem is particularly acute because we are simulating a large complex that includes multiple disordered regions interacting with each other. Despite sampling limitations, the contact probabilities reported here allow us to identify key interactions of interest, which could direct further mutation studies. Our study describes a general approach for simulations of large biomolecular systems whose interaction interface consists of highly flexible disordered regions. This approach could be adapted to study similar dynamic interfaces, which are particularly relevant in understanding the interactions between viral and host proteins.

## Materials and methods

### Protein Model Construction

We constructed a homology model for human TRIM5*α* PRYSPRY domain using both the NMR structure (PDB:2LM3^14^) and the crystal structure (PDB: 4B3N^18^) of rhesus macaque TRIM5*α*. We used Modeller^52,53^ with the first isoform in the UNIPROT entry for human TRIM5*α* (UNIPROT code Q9C035) as the target sequence. We also retrieved the human TRIM5*α* structure predicted by AlphaFold Monomer V2.0^22^ and compared its MolProbity^54^ score to that of the homology model constructed using Modeller. The structure predicted by AlphaFold compared favourably to the homology model in the MolProbity scores. For this reason, we chose to use the AlphaFold structure as the basis for our structural model of human TRIM5*α*. We included only the residues belonging to the PRYSPRY domain (residues 290-493, inclusive). PyMOL^55^ was used to introduce the R332P mutation. Because AlphaFold predicts the structure of disordered regions with low confidence,^41^ we remodelled the disordered V1 loop (residue 324-345) using Modeller.^52,53^ Loop refinement was performed separately on huTRIM5*α* and huTRIM5*α*^R332P^. For each system, 1000 loop models were generated for residues 324-345, and the 9 with the lowest Discrete Optimized Protein Energy (DOPE) score^56^ were chosen as the starting structures for MD simulations. The initial structure for rhTRIM5*α* was constructed by introducing the mutation T307P in the NMR structure for rhTRIM5*α* (PDB 2LM3^14^) using PyMOL to recover the wild-type sequence.

To construct the initial conformation for the HIV capsid-rhTRIM5*α* complex simulations, we started with the cryo-electron microscopy structure of the HIV capsid (PDB: 3J3Y).^25^ We extracted a trimer of hexamers of the capsid protein from this structure. Specifically, we chose the trimer defined by atom IDs 19812-30616 (bundle 3), 55832-66636 (bundle 4) and 77444-88249 (bundle 4). The choice of which trimer of hexamers to use was arbitrary. We used structures from the coarse-grained model of Yu et al.^23^ as a basis for the relative orientation of the HIV capsid trimer-TRIM5*α* complex. We extracted an arbitrary capsid trimer-PRYSPRY domain complex from the coarse-grained structure file. The initial structure of the rhTRIM5*α* PRYSPRY domain was taken from our simulations. In order to ensure that we picked a representative conformation of the PRYSPRY domain to construct the complex, we first characterized its conformational landscape. To do this, we performed PCA on the simulated conformational ensembles of all three simulated systems (Figure S9). The feature vector we used was the pairwise C*α* distances of residues 324-349, 376-400, 414-430 (huTRIM5*α* residue numbering), excluding the two insertions in the V1 loop of rhTRIM5*α*. This selection is primarily constituted by the flexible V1-V3 loops, which together represent the most conformational variation in the system, and are known to be involved in capsid binding. In this 2D PCA space, each point represents a conformation from the simulation trajectories. We used the DBSCAN^47^ clustering algorithm to identify densely populated states in this conformational landscape (Figure 4a). The clustering analysis resulted in four clusters. One structure was randomly selected from each of the four clusters to attempt to construct the complex. We used MDAnalysis^57^ to align the all-atom HIV capsid trimer to the coarse-grained HIV capsid trimer, and the all-atom rhTRIM5*α* structures to the coarsegrained rhTRIM5*α* structures. Three out of the four rhTRIM5*α* structures led to steric clashes between the PRYSPRY domain and the HIV capsid trimer (an example is shown in Figure S10). One structure, taken from cluster 3 (Figure 4a,c), aligned without any steric clashes and was used as the starting structure for simulations of the complex.

### MD Simulations - PRYSPRY Domains of TRIM5*α* variants

We simulated PRYSPRY domains from three TRIM5*α* variants: rhTRIM5*α*, human TRIM5*α*, and huTRIM5*α*^R332P^ (Table 1). These simulations were performed using GRO-MACS^58^ 2021.4. The CHARMM36m force field^59^ was used with the CHARMM-modified TIP3P water model.^60^ The N- and C-termini were simulated in the charged state and all titratable residues were simulated in their standard states for pH 7. The protein was placed in a triclinic box with 1.3 nm distance to all box edges. Periodic boundary conditions were used. Water molecules were added to the simulation box and neutralizing Na+ and Cl- ions were added at a concentration of 150 mM. The energy of each system was minimized using the steepest descent method. Then, an equilibration simulation was carried out for 20 ns in in the NVT ensemble using the velocity-rescale temperature coupling method^61^ with position restraints on all heavy atoms (with force constants of 1000 kJ/mol). This was followed by an equilibration simulation in the NPT ensemble using the Berendsen pressure coupling method,^62^ again with position restraints on all heavy atoms with force constants of 1000 kJ/mol. Next, a 20 ns equilibration simulation was performed in the NPT ensemble with the Parrinello-Rahman pressure coupling method.^63^ The Parrinello-Rahman pressure coupling method was used for all subsequent simulations. Then, position restraints were first released, gradually, from the disordered V1 loop (324-345 in human, 326-349 in rhesus) to allow it to equilibrate before the rest of the protein. Every 20 nanoseconds, the position restraint force constants on loop residues were reduced by the following regimen: 500 kJ/mol,250 kJ/mol, 125 kJ/mol, 75 kJ/mol, 35 kJ/mol, 20 kJ/mol, 10 kJ/mol, 5 kJ/mol, 1 kJ/mol, 0 kJ/mol. Then, an identical regimen was applied to all other heavy atoms, for a grand total of 460 ns of equilibration simulation per trajectory. Production MD was then performed in the NPT ensemble using the Parrinello-Rahman barostat and the v-rescale thermostat. Nine independent trajectories of 8 *µ*s were generated for each system. A simulation timestep of 2 fs was used and hydrogen bonds were constrained using the LINCS algorithm.^64^ A cut-off of 1.2 nm was used to calculate short range electrostatic and van der Waals interactions. The verlet cutoff-scheme was used. The non-bonded pair list was updated every 40 fs to include atoms less than 1.2 nm apart. Long range electrostatic interactions were computed using Particle-mesh Ewald summation^65^ with a grid spacing of 0.16 and fourth order cubic interpolation. All simulations were performed at a temperature of 298 K and a pressure of 1 bar.

### MD Simulations - HIV Capsid-TRIM complex Systems

The simulation setup and parameters were the same as described above for the simulations of the PRYSPRY domains. The energy of the system was minimized using the steepest descent method. An equilibration simulation was carried out in the NVT ensemble for 2 ns using the velocity-rescale temperature coupling method^61^ and position restraints on all heavy atoms with a force constant of 1000 kJ/mol. Then, we carried out 100ns of simulation in the NPT ensemble with the Berendsen barostat.^62^ We extracted protein structures from time points 20ns, 40ns, 60ns, 80ns, and 100ns of this simulation to spawn 5 independent simulations. Each of these structures was placed in a dodecahedral box with 1 nm distance to all box edges. Periodic boundary conditions were used. Water molecules^60^ were added to the simulation box and neutralizing Na+ and Cl- ions were added at a concentration of 150 mM. The energy of each system was minimized using the steepest descent method. An equilibration simulation was carried out in the NVT ensemble for 2 ns using the velocity-rescale temperature coupling method^61^ and position restraints on all heavy atoms with a force constant of 1000 kJ/mol. Then, we carried out 2 ns of simulation in the NPT ensemble with the Berendsen barostat.^62^ Finally, each system was simulated in the NPT ensemble with the Parrinello-Rahman pressure coupling method^63^ for 500 ns.

### Analysis of Simulation Trajectories

Given the length of the equilibration simulation runs, all production simulation data was used for analysis. All coordinate-based trajectory analysis was performed using the MDAnalysis module version 2.2.0^57^ for Python 3, including R*_g_*, atomic distances, RMSF, and contacts. PCA was performed using the scikit-learn^66^ module for Python 3. Secondary structure was determined with the DSSP^67^ function in the MDTRAJ^68^ module for Python 3. Hydrogen bonds were determined using MDAnalysis with a 3.5 °*A* distance cutoff between donor and acceptor atoms, and 150^◦^ donor-hydrogen-acceptor angle cutoff. Two residues were considered to be in contact based on a distance cutoff of 4.5 °*A* between any pair of heavy atoms. Uncertainty is reported as the standard error of the mean, with each independent trajectory being treated as an independent measurement (unless otherwise stated). B-factors were computed from RMSF values using Equation 1.^69^

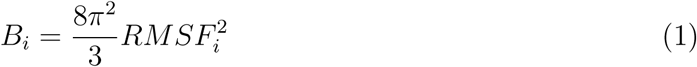

### Cell culture

Cell lines HEK293T/17 (CRL-11268) – a transfectable cell line for virus production – and CRFK (CCL-94) – a cat fibroblast cell line – were purchased from the ATCC. CRFK cells stably transduced with a human TRIM5*α* (either wild-type or an R332P variant) or a matched empty vector, each of which contained a puromycin resistance cassette, were described previously.^10^ Cells were grown on tissue-culture treated plates in DMEM containing high glucose and L-glutamine (Gibco) and supplemented with 1x penicillin/streptomycin (Gibco) and 10% fetal bovine serum (Gibco); TRIM5*α* expression in CRFK cells was maintained by selection with 2 *µ*g/mL puromycin (Fisher). Cells were grown at 37^◦^C in 5% CO2 in humidified incubators. Cells were harvested from plates by digestion with 0.05% trypsin-EDTA (Thermo Fisher) and counted using a TC20 (Biorad) automated cell counter.

### Plasmids and cloning

HIV-1 virus-like particles were generated from three plasmids: two plasmids for transient expression of the VSV-G pseudotyping envelope protein (pMD2.G, Addgene #12259) and the HIV-1 gag-pol (pSPAX2), and an HIV-1 transfer vector encoding GFP pHIV-ZsGreen.^70^ Mutations at capsid residue R82 were generated by site-directed mutagenesis of pSPAX2 using primers containing the desired mutations flanked by perfect homology on each side (∼20 nucleotides, ∼52^◦^C melting temperature): primers for R82L (fwd: ggaagctgcagaatgggatTTagtgcatccagtgcatg, rev: catgcactggatgcactAAatcccattctgcagcttcc), R82M (fwd: ggaagctgcagaatgggataTGgtgcatccagtgcatgcag, rev: ctgcatgcactggatgcacCAtatcccattctgcagcttcc), and R82K (fwd: ggaagctgcagaatgggataAagtgcatccagtgcatgcag, rev: ctgcatgcactggatgcactTtatcccattctgcagcttcc). Primestar HS polymerase (Takara) was used to minimize errors during full-plasmid amplification (98^◦^C 10”; 24 x [98^◦^C 10”;, 62^◦^C, 15”;, 72^◦^C, 8.5’]; 72^◦^C, 20’), followed by DpnI digestion of unmodified parental DNA and chemical transformation into NEB5*α* (NEB). DNA was isolated from individual colonies by PureYield miniprep (Promega) and sequenced using an HIV1gag-F primer (GCACAGCAAGCAGCAGC) to identify correct clones. All DNA used for virus production was purified by Nucleobond Xtra Midiprep kits (Takara).

### HIV-1 production and infection

Single-cycle HIV-1 virus-like particles, which are capable of only one round of infection, were generated by transfecting HEK-293T/17 cells, seeded the prior day at 5 x 10^5^ cells/mL in 6-well plates, with 3 plasmids (1 *µ*g of GFP-encoding transfer vector pHIV-ZsGreen, 667 ng of the appropriate pSPAX2 variant, and 333 ng of pMD2.G per well) using 3 *µ*L of Trans-IT 293T transfection reagent (Mirus Bio) per *µ*g of DNA. After 12-24 hours, media was aspirated and replaced with 1 mL of fresh media. Virus-containing supernatants were harvested at 48 hr post-transfection, clarified at 500 x g for 5 min to remove cell debris, and snap frozen in liquid nitrogen for storage at -80^◦^C.

To assess TRIM5*α*-mediated restriction, CRFK cells expressing empty vector, wild-type human TRIM5*α*, or human TRIM5*α*-R332P were seeded at 1 x 10^5^ cells/mL with 100 *µ*L/well in 96-well plates the day prior to transduction. Freshly thawed viruses were serially diluted in media containing 20 *µ*g/mL DEAE-dextran. CRFK media was aspirated and replaced with diluted virus at ½ x volume (i.e. 50 *µ*L/well). Plates were centrifuged at 1100 x g for 30 min at 30^◦^C. The following day, virus was removed and fed fresh media. Infection was monitored by GFP positivity using flow cytometry at 72 hrs post infection. Cells were gated on size (FSC vs. SSC), then single cells (FSC height vs. area), and then GFP+ or HA+ (as compared to uninfected controls; FITC vs. PE).

Virus inhibition was calculated by comparing infection of CRFK cells expressing empty vector (pQCXIP) or TRIM5*α* in the same experiment. Wells with *<* 0.5% or *>* 50% GFP+ cells were excluded due to noise or saturation, respectively. A linear regression (against logtransformed data) was then used to calculate the viral dose corresponding to 5% infection (ID5) for each cell line. The ID5 for cells expressing TRIM5*α* was normalized to empty-vector-expressing cells to yield fold restriction. All fold inhibitions were calculated from at least three independent experiments.

## Acknowledgments

This research was supported by the Research and Scholarly Activity Fund of the University of Toronto Mississauga Office of the Vice-Principal Research & Innovation and a Natural Science and Engineering Research Council of Canada (NSERC) Discovery Grant (RGPIN-201806408). This research was enabled in part by support provided by Calcul Quebec (www.calculquebec.ca) and the Digital Research Alliance of Canada (alliance.can.ca), as well as an HHMI Hanna H. Gray Fellowship (GT11096/GT16732 to JLT).

## Data Availability

MD simulation data (initial coordinates and simulation trajectories) are provided for all systems as a Zenodo repository accessible here: https://doi.org/10.5281/zenodo.14014166 Data underlying figures are also available in this Zenodo repository.

