## Supplementary Information for "An all-atom view into the disordered interaction interface of the TRIM5*α* PRYSPRY domain and the HIV capsid"

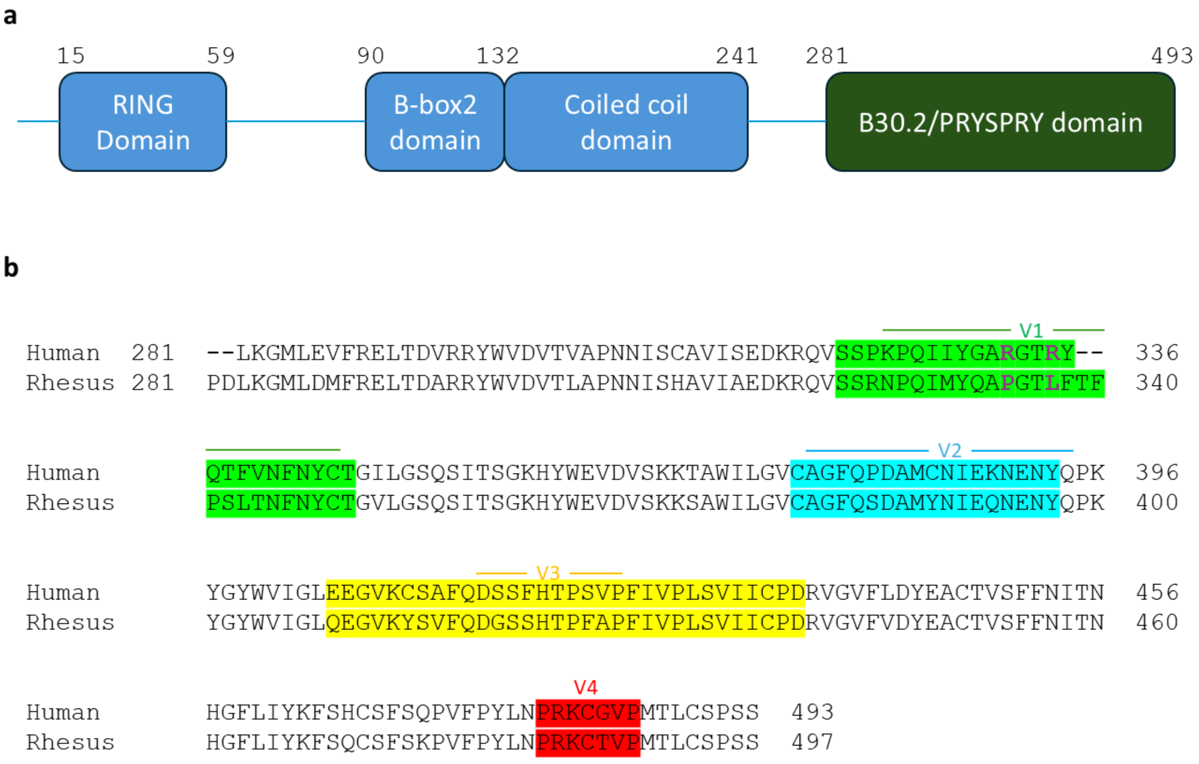

Figure 1: **Sequence and domains of TRIM5$\alpha$.** (a) Domain organization of TRIM5$\alpha$. Residue numbers shown correspond to the sequence of huTRIM5$\alpha$, with domain boundaries defined as in Tenthorey *et al.*[1] (b) Sequence alignment of huTRIM5$\alpha$ and rhTRIM5$\alpha$ PRYSPRY domains. Clustal Omega[2] was used to obtain the alignment. The two sequences are 83% identical (180/217 residues) and 88% similar (191/217 residues). Residues R332/P334 and R335/L337 (in human/rhesus numbering) are indicated in bold magenta letters. The V1-V4 variable regions (as defined by Yang *et al.*[3]) are indicated by shading on the alignment (V1 region: 323-350, V2 region: 380-397, V3 region: 409-440, V4 region: 483-489 in rhesus numbering). The V1, V2 and V3 loops are indicated by lines above the alignment (V1 loop: 326-349, V2 loop: 381-398, V3 loop: 419-428 in rhesus numbering). The V1 loop was defined consistently with previous literature.[4] The V2 and V3 loops were defined based on the secondary structure propensities in the simulations (Figure S2).

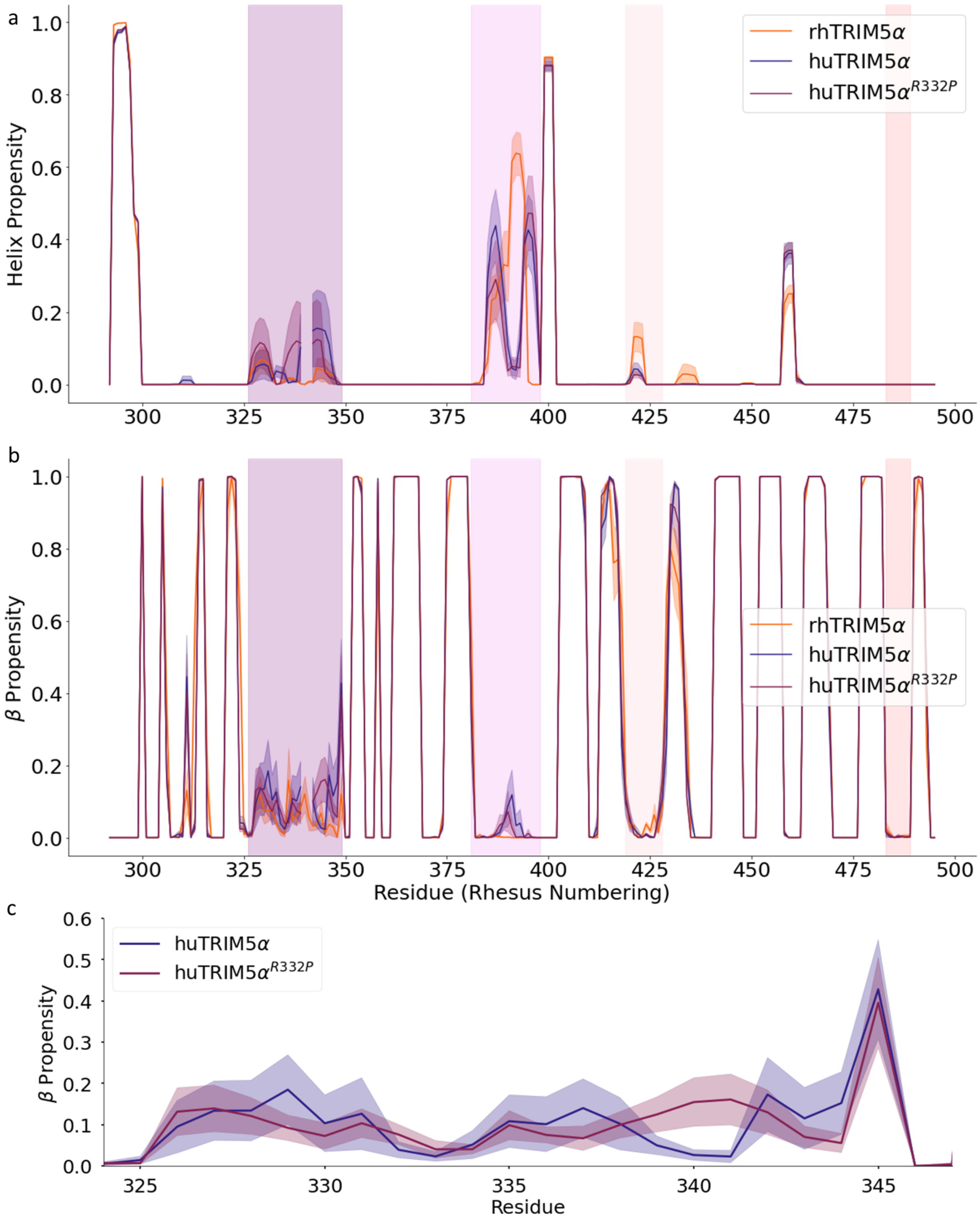

Figure 2: **Secondary structure propensity of TRIM5$\alpha$.** (a) Helix propensity for rhTRIM5$\alpha$, huTRIM5$\alpha$, and huTRIM5$\alpha^{\text{R332P}}$ computed from the simulation ensembles. (b) $\beta$ propensity for rhTRIM5$\alpha$, huTRIM5$\alpha$, and huTRIM5$\alpha^{\text{R332P}}$ computed from the simulation ensembles. In (a) and (b), vertical shaded areas indicate the V1-V4 loops. (c) $\beta$ propensity for the V1 loop of huTRIM5$\alpha$, and huTRIM5$\alpha^{\text{R332P}}$. Secondary structure was computed using DSSP[5] for each frame in each trajectory. The average helix or $\beta$ propensity is defined as the portion of frames that each residue was in an $\alpha$-helix or $\beta$-sheet conformation, respectively. Shaded regions indicate standard error of the mean obtained by treating each trajectory as an independent measurement.

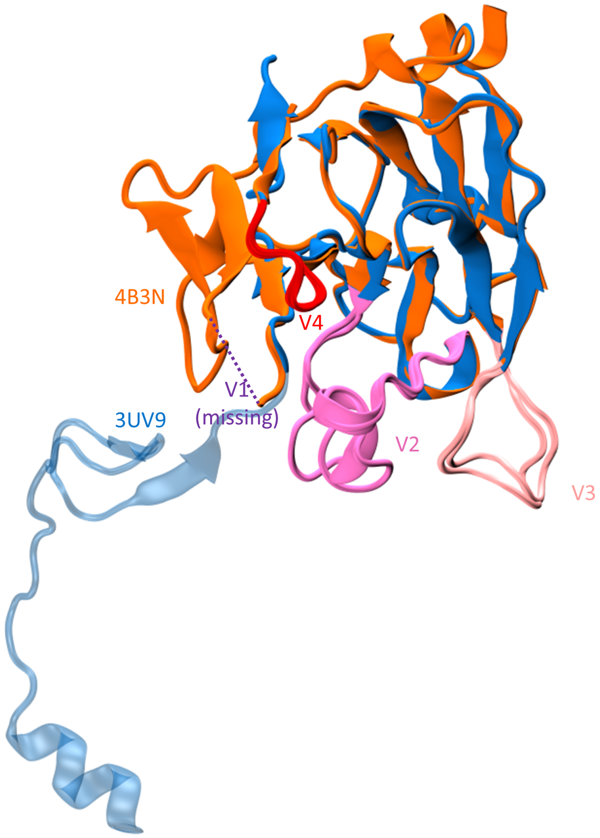

Figure 3: **Crystal Structures of rhTRIM5$\alpha$.** Two crystal structures of rhTRIM5$\alpha$ are shown (PDB ID: 4B3N[3] in orange and PDB ID: 3UV9[6] in dark blue). The V2, V3, and V4 loops are shown in mauve, pink, and red, respectively, for both structures. In the crystal structure 4B3N, the V1 loop was included but electron density was not observed. The location of the missing V1 loop in 4B3N is indicated (purple). In the crystal structure 3UV9, the V1 loop is replaced by a two residue linker, and the protein crystallized as a domain-swapped trimer in which the part of the sequence preceding the V1 loop forms contacts with neighbouring proteins in the crystal. The domain-swapped segment of 3UV9 is shown in light blue as translucent.

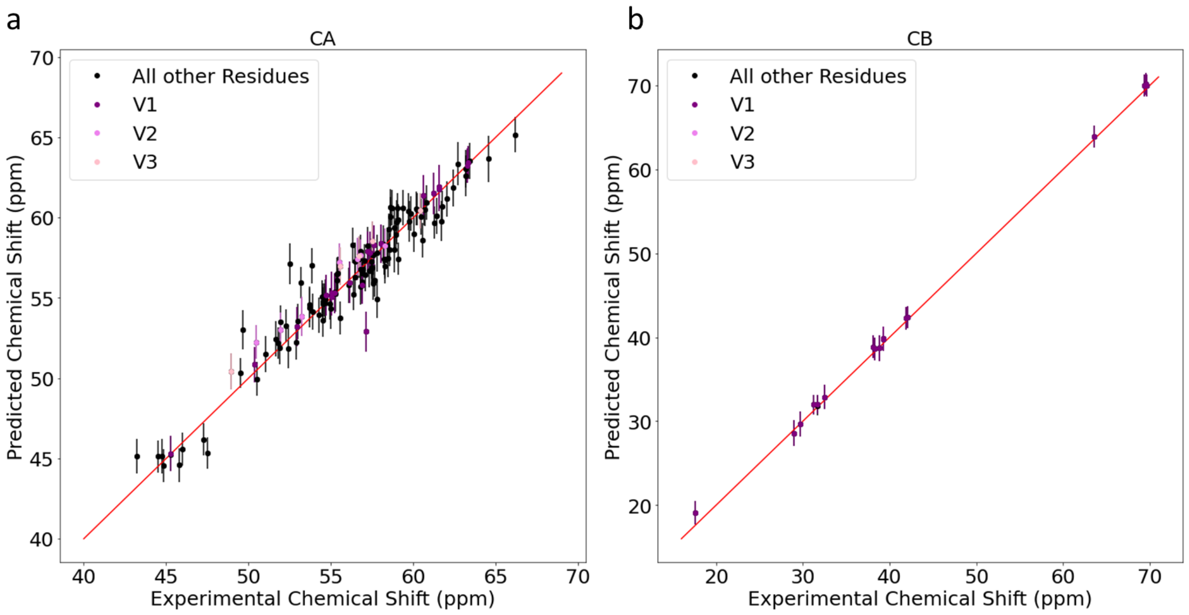

Figure 4: **Comparison of predicted and experimental chemical shifts.** A comparison of predicted chemical shifts from the simulation ensembles (obtained using SHIFTX+[7]) versus the experimental chemical shifts of rhTRIM5$\alpha$ (Biological Magnetic Resonance Bank, BMRB ID: 180978[8]). (a) C$\alpha$ chemical shifts. The average unsigned error is 0.82 $\pm$ 0.91 ppm. (b) C$\beta$ chemical shifts. The average unsigned error is 0.47 $\pm$ 0.94 ppm. For both (a) and (b), error bars indicate the standard error of the mean obtained by treating each trajectory as an independent measurement plus the RMS error of SHIFTX+ for C$\alpha$ (0.8743 ppm) and C$\beta$ (1.0099 ppm) chemical shifts.[7]

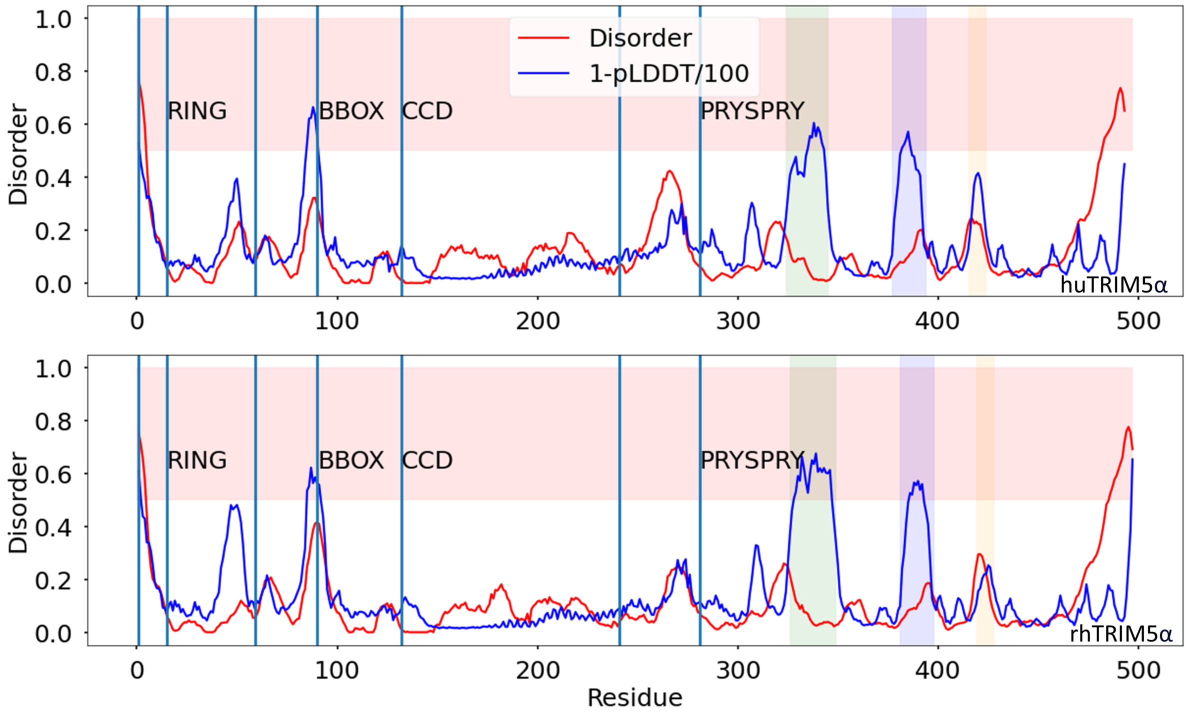

Figure 5: **Predicted disorder in TRIM5$\alpha$.** The predicted disorder by residue of huTRIM5$\alpha$ (upper plot) and rhTRIM5$\alpha$ (lower plot), as predicted by Metapredict Version 2[9,10] (red line). Higher values indicate a higher likelihood of the residue being disordered. Above a threshold value of 0.5, the residue is predicted to be disordered,[9,10] which is indicated by light red shading. The AlphaFold[11] pLDDT score is normalized and sign-reversed (1-pLDDT/100). It is shown as a blue line to allow comparison to the Metapredict Version 2 prediction. Domain boundaries are as defined by Tenthorey *et al.*[1] and are indicated by vertical lines. The loops are indicated by shading (V1 loop, green; V2 loop, blue; V3 loop, yellow).

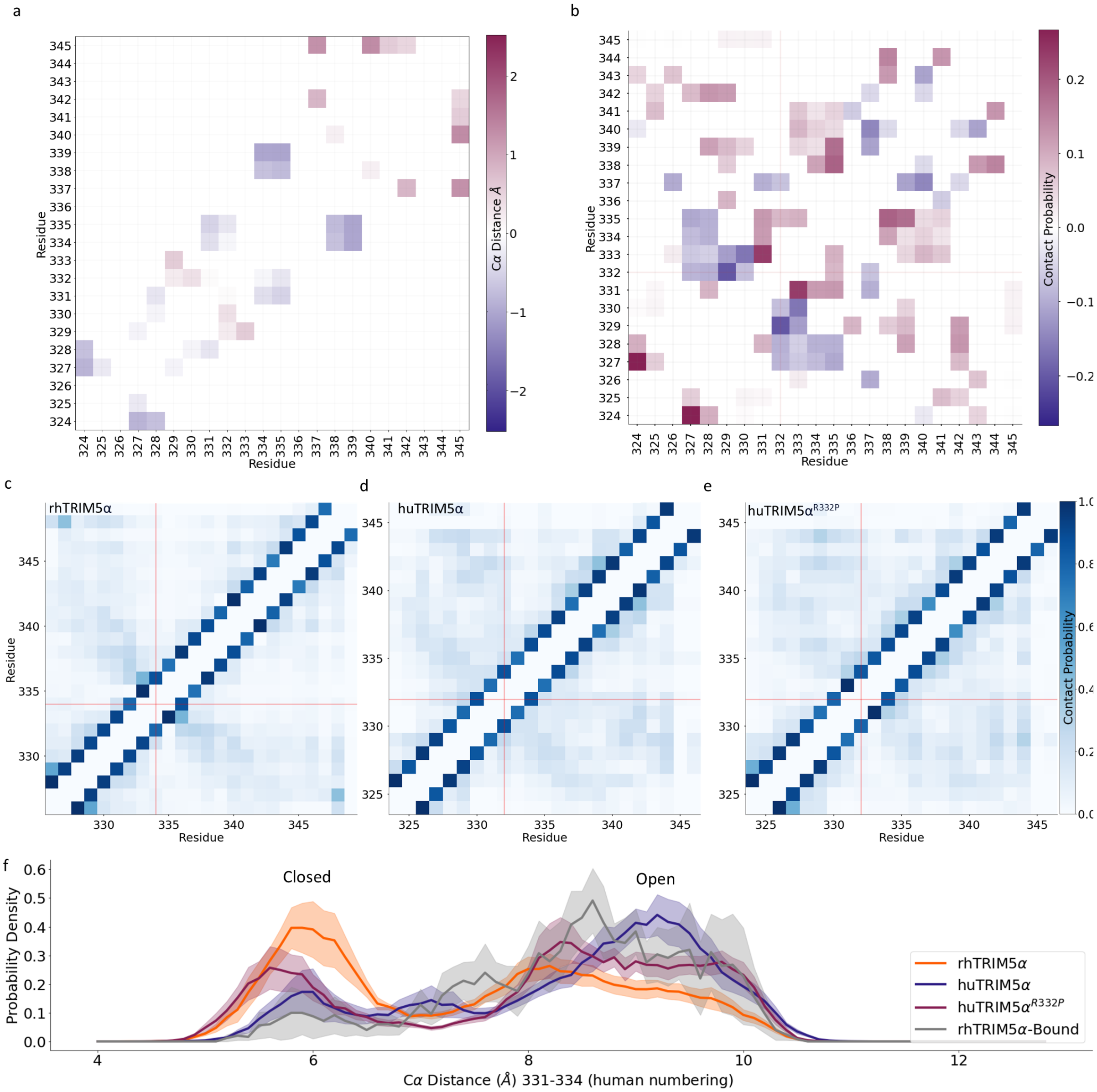

Figure 6: **Structural differences between TRIM5$\alpha$ variants in the V1 loop.**. (a) Difference map of C$\alpha$ distances (the average C$\alpha$ distance map of huTRIM5$\alpha^{\mathrm{R332P}}$ minus that of huTRIM5$\alpha$). Only statistically significant differences are shown. Distances that are larger in huTRIM5$\alpha^{\mathrm{R332P}}$ are more plum coloured and distances that are larger in huTRIM5$\alpha$ are more blue. (b) The contact difference map for huTRIM5$\alpha^{R332P}$ minus huTRIM5$\alpha$ for the residues in the V1 loop only. Contact probabilities were computed for each system and the difference was taken. Plum indicates that a contact is formed more often in huTRIM5$\alpha^{R332P}$, while blue indicates that a contact is formed more often in huTRIM5$\alpha$. (c-e) The contact maps for residues in the V1 loop for rhTRIM5$\alpha$ (c), huTRIM5$\alpha$ (d), and huTRIM5$\alpha^{\mathrm{R332P}}$ (e). The coloring of each square in the contact maps in (c-e) indicates the probability of a given contact. (f) Probability distribution of the distance between C$\alpha$ atoms of residues 331 and 334 (corresponding to residues 333 and 336 in rhTRIM5$\alpha$) from the simulations. rhTRIM5$\alpha$ is shown in orange, huTRIM5$\alpha$ is shown in navy, and huTRIM5$\alpha^{\mathrm{R332P}}$ is shown in plum, rhTRIM5$\alpha$ in the complex with the capsid is shown in grey. Shading indicates standard error of the mean obtained by treating each trajectory as independent.

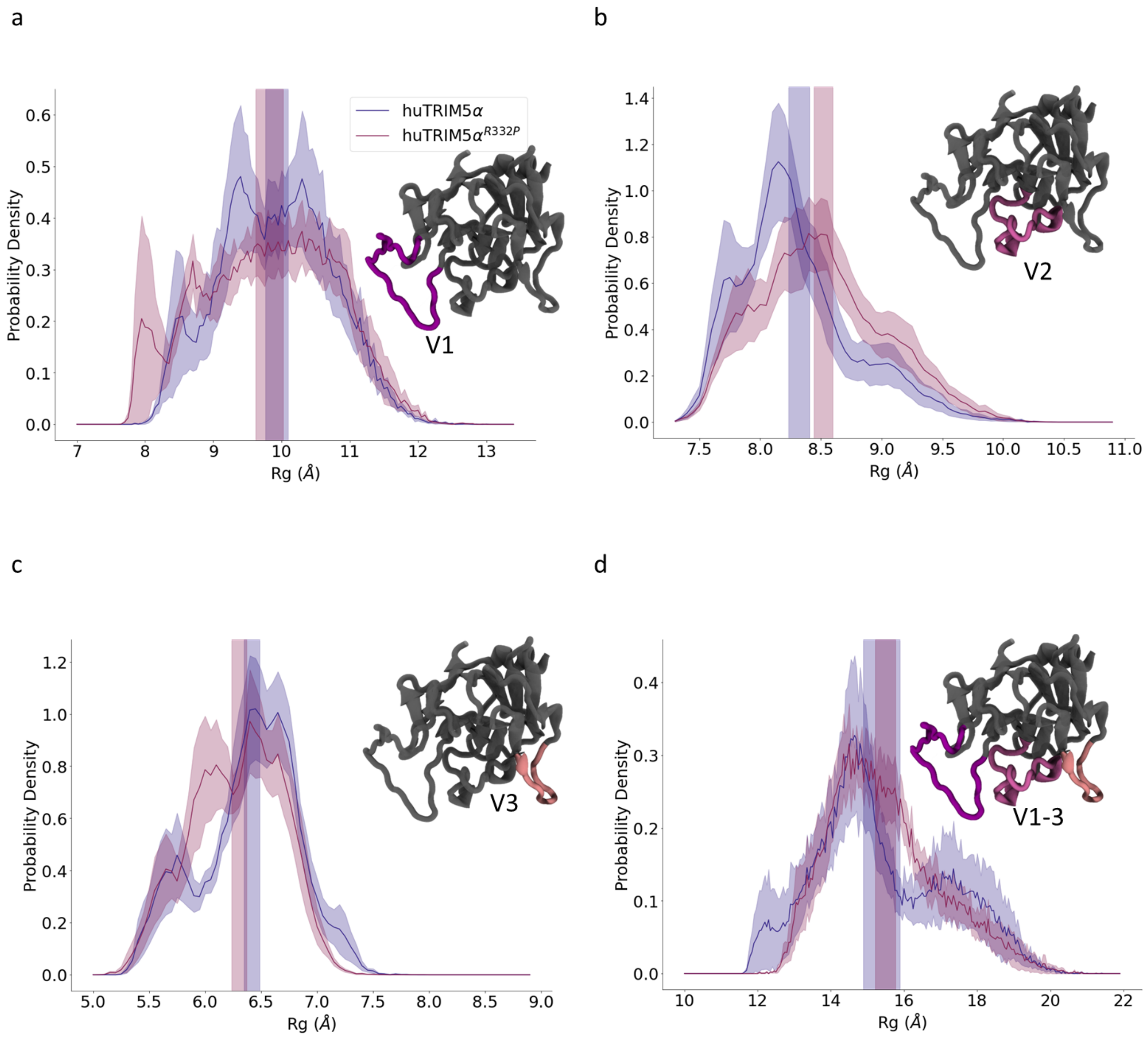

Figure 7: **Analysis of the R$_g$ of huTRIM5$\alpha$ loops.** (a) R$_g$ distribution of the V1 loop. (b) R$_g$ distribution of the V2 loop. (c) R$_g$ distribution of the V3 loop. (d) R$_g$ distribution of the V1-V3 loops (including all residues in each of the three loops). huTRIM5$\alpha$ is shown in navy and huTRIM5$\alpha^{\text{R332P}}$ is shown in plum. Shaded regions indicate standard error of the mean obtained by treating each trajectory as independent. 3D structures are taken from the simulations of huTRIM5$\alpha$.

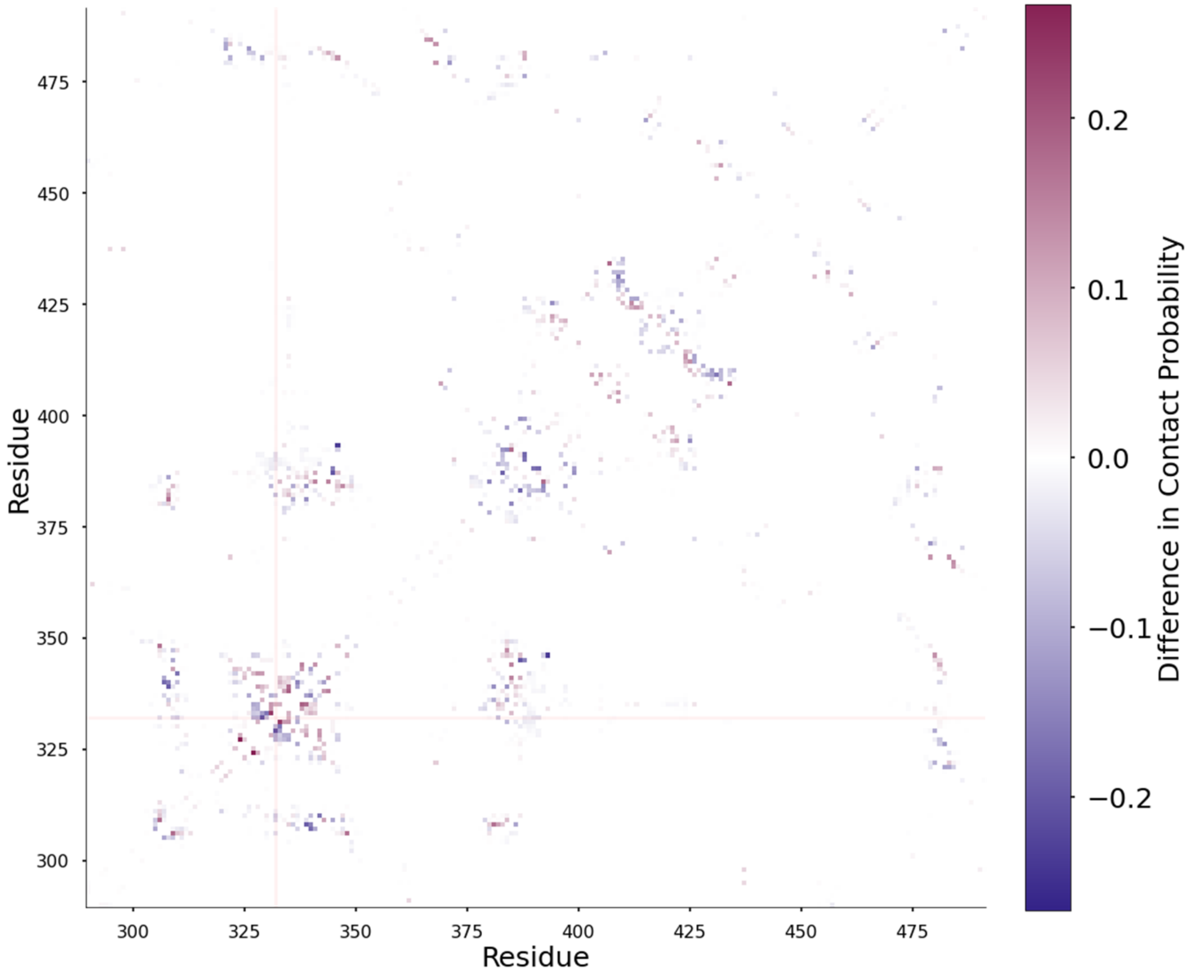

Figure 8: **Contact difference map for huTRIM5$\alpha^{R332P}$ minus huTRIM5$\alpha$.** The contact probability difference map for huTRIM5$\alpha^{R332P}$ minus huTRIM5$\alpha$. Contact probabilities were computed for each system and the difference was taken. Plum indicates that a contact is formed more often in huTRIM5$\alpha^{R332P}$, while blue indicates that a contact is formed more often in huTRIM5$\alpha$. A close up of the contact difference map for the residues in the V1 loop is shown in Figure S6b. For reference, the residue numbers of the loops are as follows, V1 loop: 324-345, V2 loop: 377-394, V3 loop: 415-424, V4: 379-385.

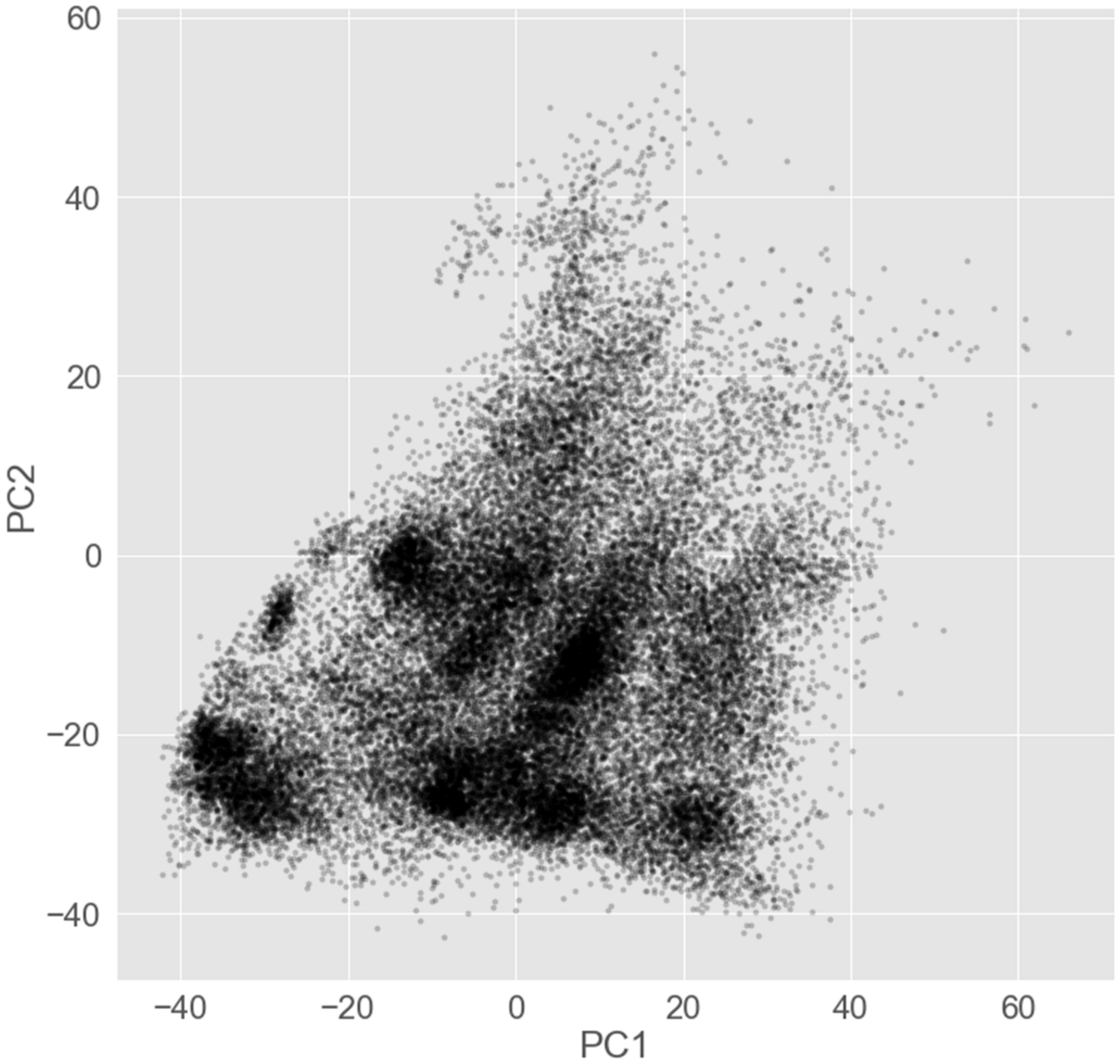

Figure 9: **Conformational landscape of the rhTRIM5$\alpha$ loops.** PCA of the rhTRIM5$\alpha$ PRYSPRY domain ensemble. Each point in the 2D space defined by the first two principal components (PC1 and PC2) corresponds to a structure from the rhTRIM5$\alpha$ PRYSPRY domain simulations (see Methods for description of the PCA).

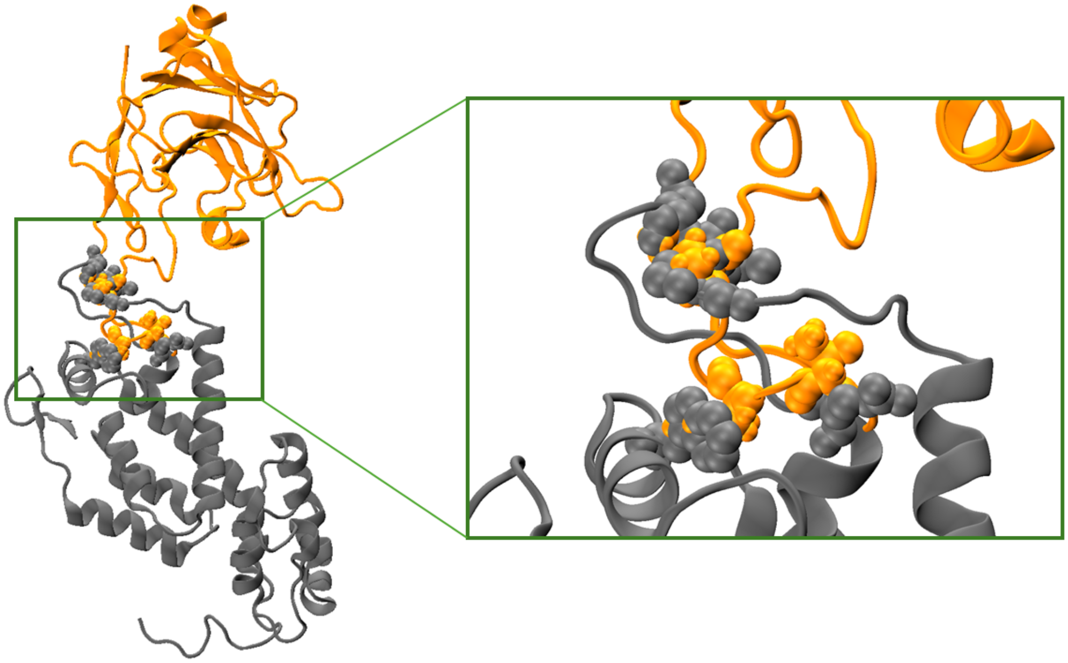

Figure 10: **Steric clashes in the aligned complex.** An example of steric clashes between loops of the aligned complex of rhTRIM5$\alpha$ and the capsid. The rhTRIM5$\alpha$ PRYSPRY domain and the capsid protein are shown in orange and gray, respectively. Residues containing atoms that have distances less than 0.5$\mathring{A}$ to atoms in the other protein are shown in van der Waals representation.

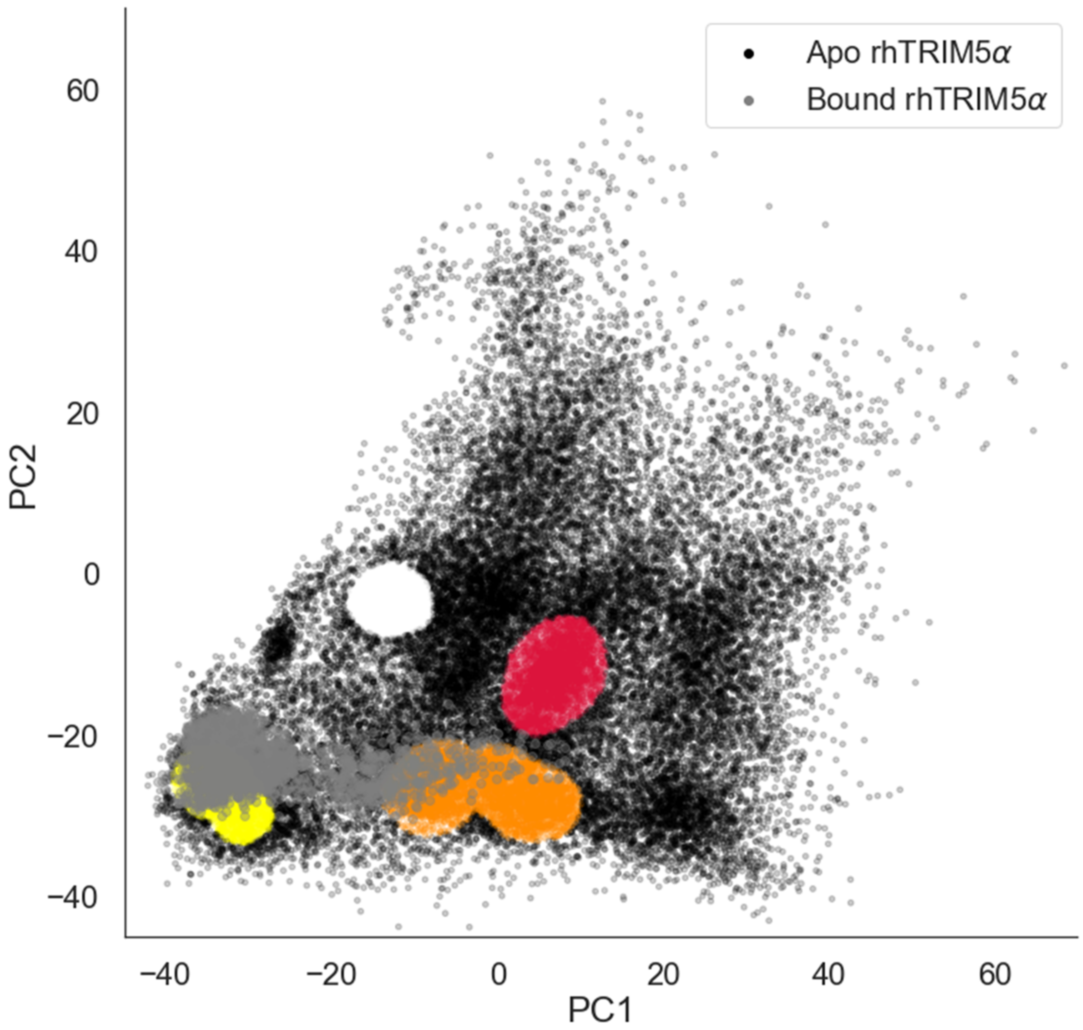

Figure 11: **Conformational landscape of rhTRIM5$\alpha$ in complex with the HIV capsid.** The conformational space of the V1-V3 loops of unbound rhTRIM5$\alpha$ is shown (as in Figure 4a). The four clusters are shown in red (1), orange (2), yellow (3), and white (4), and the outliers are shown in black. Structures from the capsid-bound rhTRIM5$\alpha$ simulations are projected onto this space (gray points).

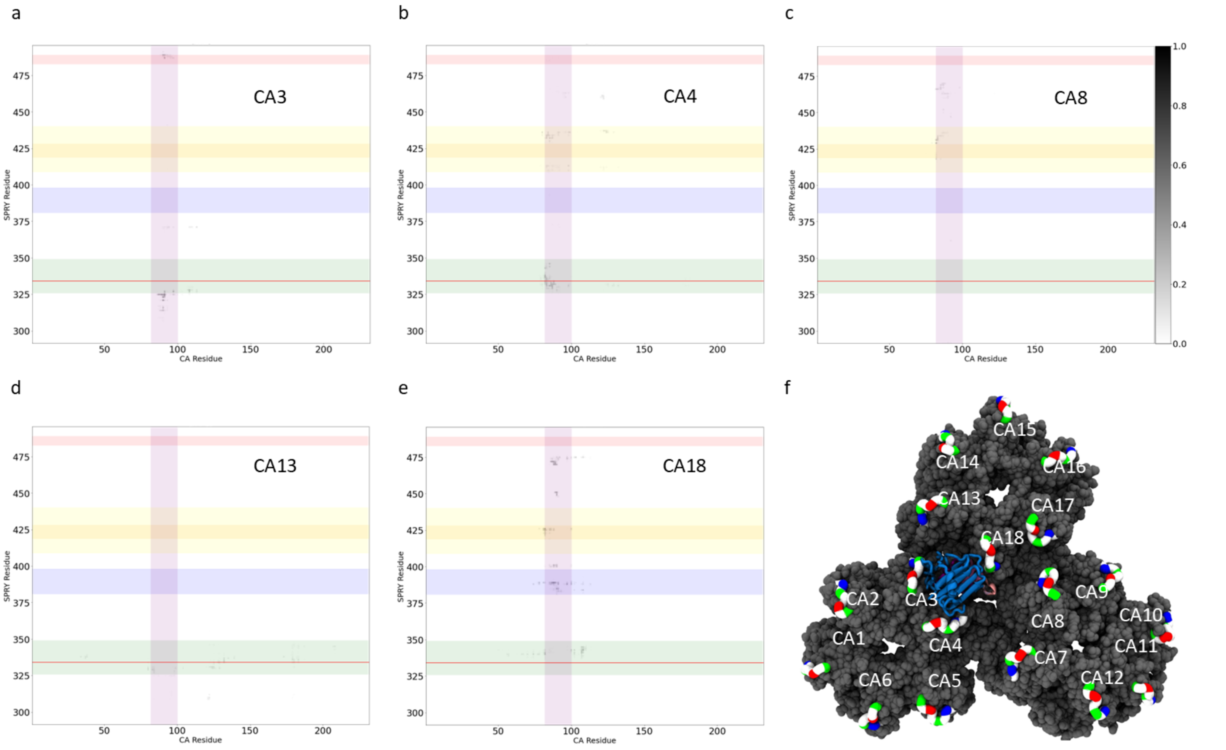

Figure 12: **Contacts between the PRYSPRY domain and individual capsid monomers.** (a-e) Probability of contacts between residues of the PRYSPRY domain and residues of the capsid protein. Each plot is labeled by the number of the capsid monomer. (f) The structure of the trimer of hexamers of the HIV capsid protein with the numbering system used for the capsid monomers. The colour scheme is the same as in Figure 4b. Note that panel (f) shows a single structure from the simulation, while the contact probability plots (a-e) are shown for all capsid monomers with which TRIM forms contacts during the five independent simulations. The majority of the contacts are present with very low probability, indicated by light gray shading.

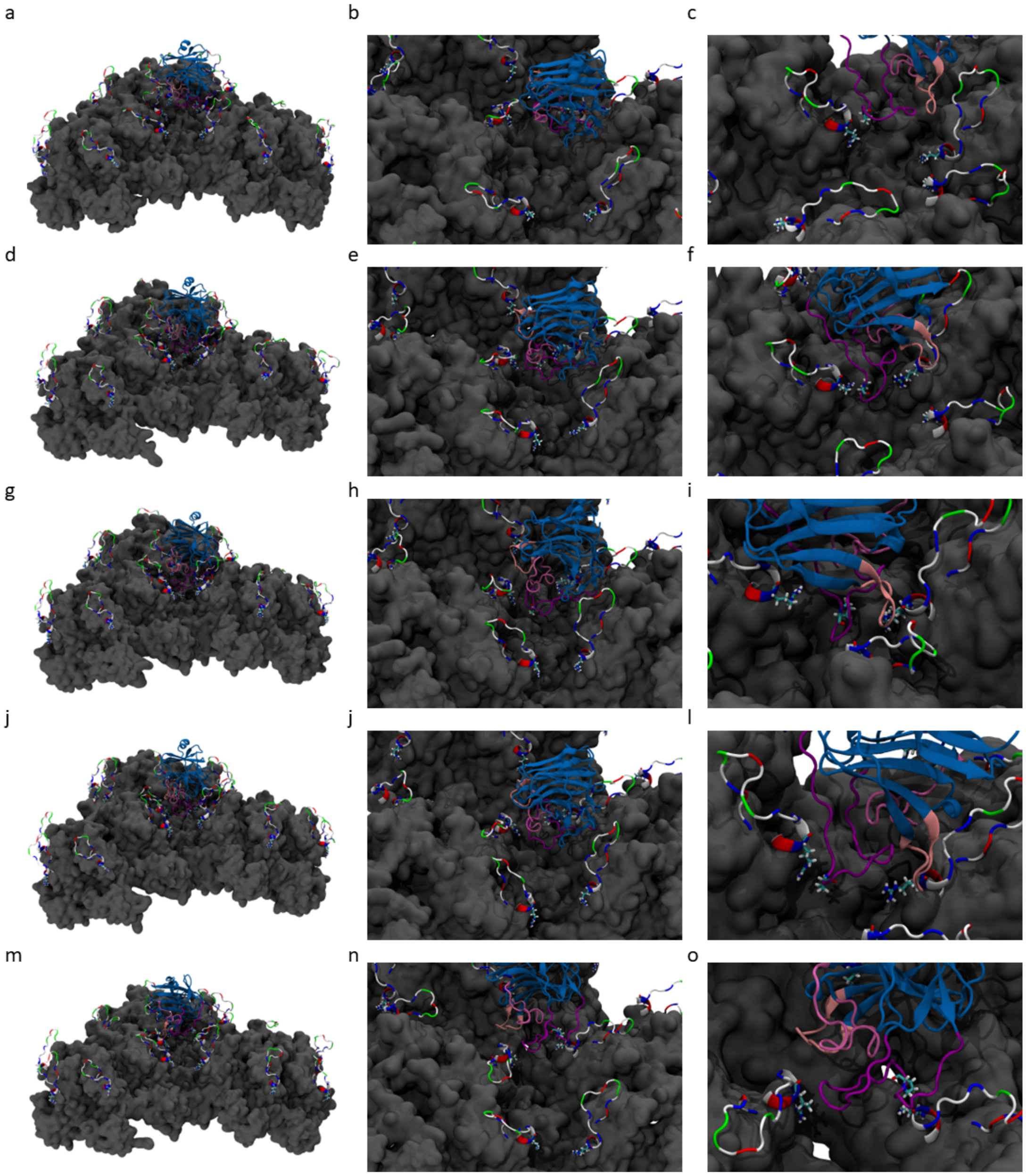

Figure 13: **Several conformations of the rhTRIM5$\alpha$ protein in complex with the HIV capsid.** Five different conformations of the HIV capsid - rhTRIM5$\alpha$ complex are shown. Each row shows a different conformation from three viewpoint angles. The first two columns show the same viewpoint angle for each conformation, and the third column shows a different angle for each conformation to improve clarity.

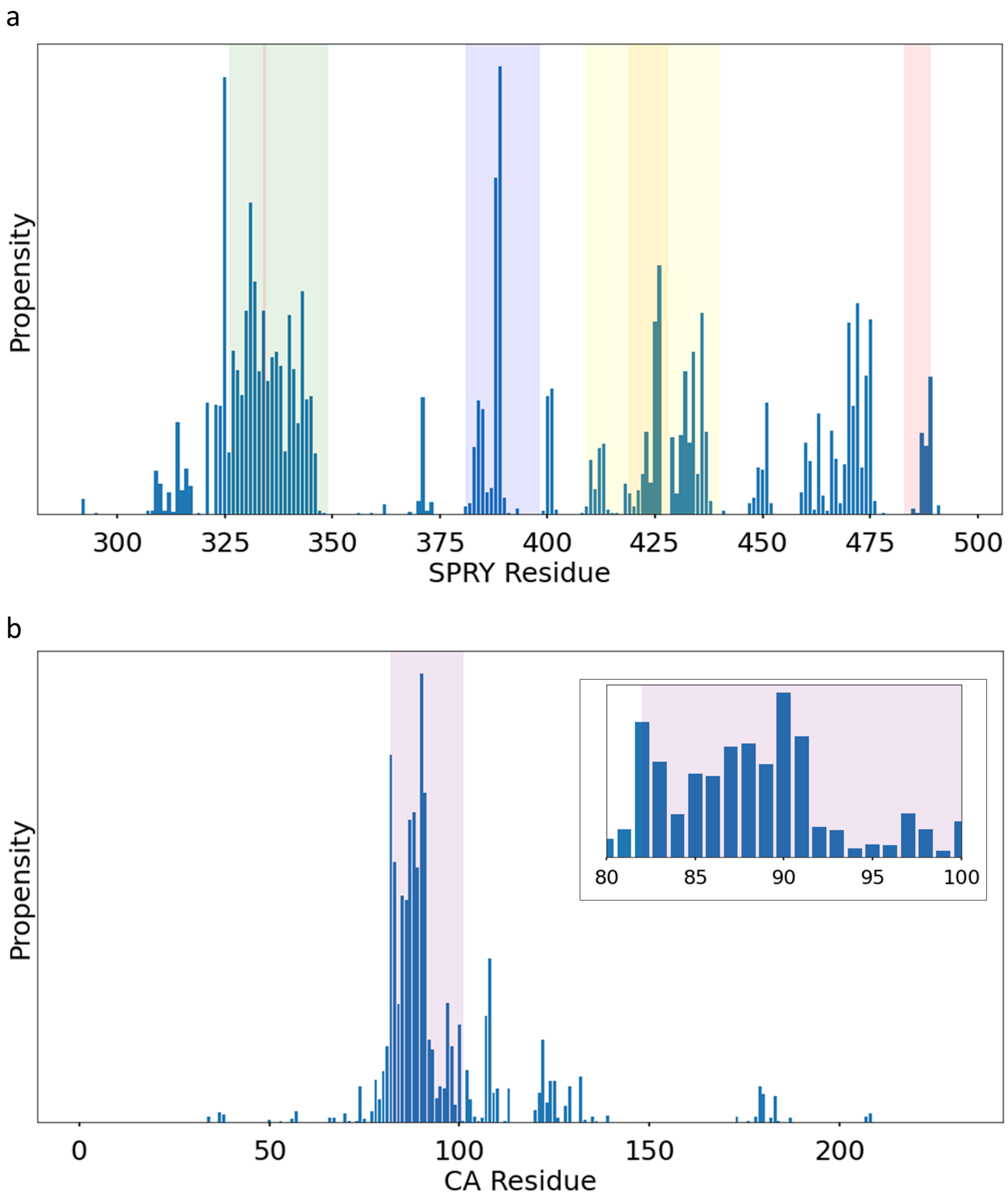

Figure 14: **Contact propensity by residue between rhTRIM5$\alpha$ and the HIV Capsid.** (a) Average relative propensity of contacts formed by the rhTRIM5$\alpha$ PRYSPRY domain. V1-V4 regions are highlighted. Shaded regions delineate various regions of each protein: green indicates the V1 loop (326-349), blue indicates the V2 loop (381-398), orange indicates the V3 loop (419-428), yellow indicates the broader V3 variable region, and red indicates the V4 loop (483-489). The light red vertical line indicates the location of residue P334 (corresponding to R332 in huTRIM5$\alpha$). (b) Average relative propensity of contacts formed by the HIV capsid protein. Residues belonging to the loop connecting helices 4 and 5 (residues 82-100), which includes the region defined by Gamble et al. as the CYPA binding loop (residues 86-95),[12] are highlighted in purple and shown in the inset for clarity.

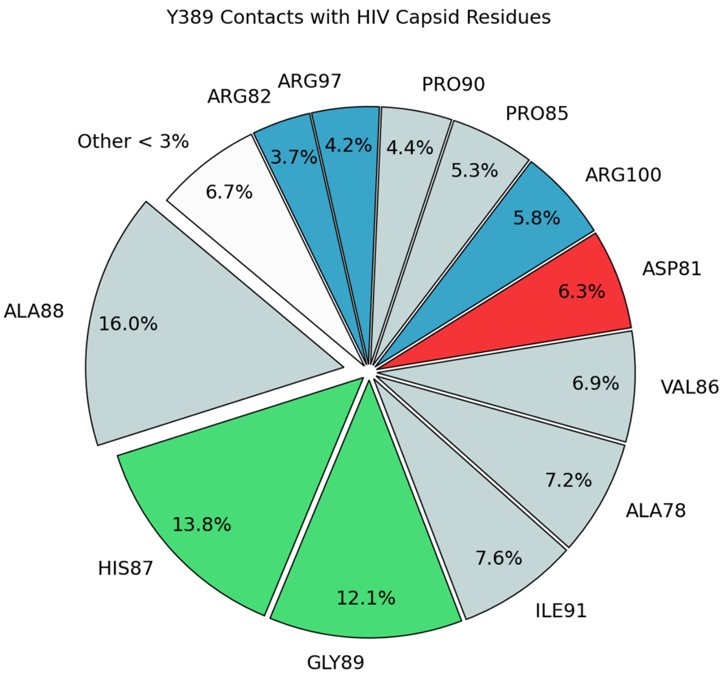

Figure 15: **Contacts formed between rhTRIM5$\alpha$ residue Y389 and the HIV capsid.** All HIV capsid protein residues that Y389 from rhTRIM5$\alpha$ forms contacts with are shown according to their respective portion of all Y389-HIV capsid contacts.
